# The molecular mechanism and physiological role of silent nociceptor activation

**DOI:** 10.1101/2022.04.07.486730

**Authors:** Timo A. Nees, Na Wang, Pavel Adamek, Clement Verkest, Carmen La Porta, Irina Schaefer, Julie Virnich, Selin Balkaya, Vincenzo Prato, Chiara Morelli, Nadja Zeitzschel, Valerie Begay, Young Jae Lee, Anke Tappe-Theodor, Gary R. Lewin, Paul A. Heppenstall, Francisco J. Taberner, Stefan G. Lechner

## Abstract

Silent nociceptors are sensory afferents that are insensitive to noxious mechanical stimuli under normal conditions but become sensitized to such stimuli during inflammation. Using RNA-sequencing and quantitative RT-PCR we demonstrate that inflammation selectively upregulates the expression of the transmembrane protein TMEM100 in silent nociceptors and electrophysiology revealed that over-expression of TMEM100 is required and sufficient to un-silence silent nociceptors. Moreover, we show that mice lacking TMEM100 do not develop secondary allodynia – i.e. pain hypersensitivity that spreads beyond the site of inflammation – in a mouse model of knee joint inflammation and that AAV-mediated overexpression of TMEM100 in articular afferents in the absence of inflammation is sufficient to induce allodynia in remote skin regions without causing knee joint pain. Thus, our work identifies TMEM100 as a key regulator of silent nociceptor un-silencing and reveals a physiological role for this hitherto enigmatic afferent subclass in triggering spatially remote secondary allodynia during inflammation.

## Introduction

Pain is an unpleasant and multifaceted sensation that can be stabbing, burning, throbbing or prickling. Likewise, pain hypersensitivity has many faces and can manifest as increased sensitivity to heat, cold or mechanical stimuli that commonly spreads far beyond the site of initial insult, a phenomenon known as secondary hyperalgesia or allodynia. Our ability to distinguish this plethora of painful sensations, relies on the functional diversity of primary sensory afferents, which detect painful stimuli and relay information about the intensity and quality of these stimuli to the central nervous system. Functionally distinct subclasses of sensory afferents have already been discovered several decades ago by classical neurophysiological studies ^1^ and, more recently, single-cell RNA-sequencing studies have mapped transcriptional signatures to these functionally classified neurons ^2,3^ and revealed changes therein associated with chronic pain ^4,5^. Moreover, knock-out, cell ablation and optogenetic studies have deciphered the contribution of various afferent subclasses to different forms of pain, such as acute pain evoked by pinprick, pinch and punctate mechanical stimuli as well as mechanical allodynia and cold allodynia associated with nerve injury ^6–10^.

Despite the clear picture regarding the contribution of different primary afferent subtypes to various forms of pain that has emerged during the past decade, the function of one of the largest subpopulations of nociceptors – the so-called ‘silent’ nociceptors – still remains enigmatic. The term ‘silent’ nociceptor denotes sensory afferents that do not respond to noxious mechanical stimuli under normal conditions but become sensitized to such stimuli during experimentally induced inflammation. Silent nociceptors were first documented in the articular nerves of the cat knee joint ^11^ and were subsequently found in the colon ^12,13^ and the bladder ^14^ as well as in human skin ^15^. Since the sensitivity of silent nociceptors to non-mechanical stimuli has never been systematically tested, it is, however, more appropriate to use the term mechanically insensitive afferents (MIAs) to describe this peculiar afferent subclass. It is estimated that MIAs constitute ∼30% of all C-fiber afferents in viscera and joints and about 15-20% in the human skin, whereas they appear to be less abundant in mouse skin ^16,17^. Considering the large proportion of MIAs in viscera and joints, it has been hypothesized that the un-silencing of MIAs during inflammation substantially increases nociceptive input onto the spinal cord, which supposedly potentiates central pain processing and eventually results in increased pain sensitivity. Microneurography from cutaneous human afferents, on the other hand, suggested that silent nociceptors might induce secondary mechanical hyperalgesia ^18^. Owing to the lack of tools that would allow the unequivocal identification or the selective genetic manipulation of MIAs, neither the mechanism underlying the un-silencing of MIAs nor their exact role in pain signaling have hitherto been deciphered.

We had previously shown that mouse MIAs express the nicotinic acetylcholine receptor alpha-3 subunit (CHRNA3) and can thus readily be identified in Tg(Chrna3-EGFP)BZ135Gsat reporter mice ^19^. Most importantly, we had shown that CHRNA3-EGFP^+^ MIAs acquire mechanosensitivity upon treatment with the inflammatory mediator nerve growth factor (NGF) and demonstrated that this process requires de-novo gene transcription. We thus here set out to identify transcriptional changes that underlie the un-silencing of CHRNA3-EGFP^+^ MIAs and to eventually utilize these findings to examine the contribution of MIAs to the generation of inflammatory pain.

## Results

### NGF treatment selectively upregulates TMEM100 in silent nociceptors

To identify proteins required for the NGF-induced acquisition of mechanosensitivity of CHRNA3-EGFP^+^ MIAs ^19^, we compared the transcriptomes of CHRNA3-EGFP^+^ neurons, cultured in the absence or presence of NGF (50 ng/ml) for 24 h, using paired-end RNAseq (Fig. 1a). This comparison showed that neither the mechanically-gated ion channel PIEZO2, which is required for mechanotransduction in CHRNA3-EGFP^+^ neurons ^19^, nor any of the known PIEZO2 modulators, such as *Stoml3, Pcnt, Mtmr2, Tmem150c, Cdh1, Anxa6, Atp2a2* and *Nedd4-2* ^20–27^, are up-regulated by NGF (Fig. 1b). Moreover, the analysis showed that CHRNA3-EGFP^+^ MIAs have a transcriptional signature – i.e. co-expression of *Ntrk1, Calca, Tac1, Trpv1, Nos1, Ly6e* and *Htr3a* but not *Cyp2j12, Prrx2* and *Etv1* (Fig. 1c) – that was previously observed in a subset of peptidergic nociceptors that were classified as PSPEP2 neurons in a large scale single-cell RNA-sequencing study ^2^. Most importantly, the RNAseq screen revealed the NGF-induced up-regulation of the transmembrane protein TMEM100 (fold-change=3.805, P=4.12E-5, N=3 samples per condition, Fig. 1b), which attracted our attention because TMEM100 was previously shown to potentiate the activity of TRPA1 ^28^ – an ion channel that plays an important role in pain signaling ^29^ – by releasing it from the inhibition by TRPV1 and both channels are expressed at significant levels in CHRNA3-EGFP^+^ neurons (Fig. 1c, d). Importantly, quantitative real-time PCR (qPCR) confirmed the up-regulation of TMEM100 in CHRNA3-EGFP^+^ MIAs and further showed that no other major nociceptor subpopulation exhibits significant changes in TMEM100 expression upon NGF treatment (Fig. 1e).

**Fig. 1.**
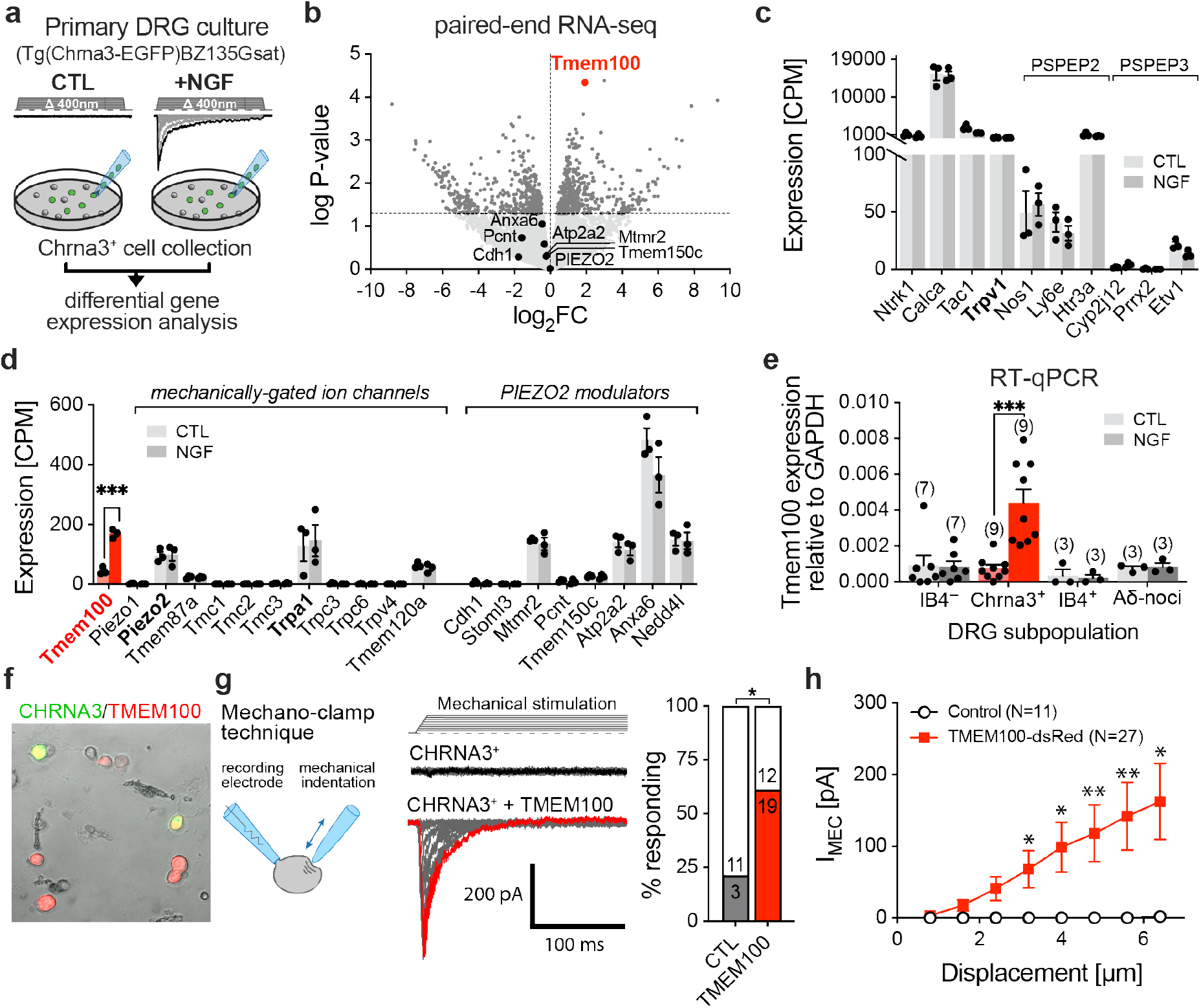
NGF treatment selectively upregulates TMEM100 in silent nociceptors. **a**, Cartoon depicting the experimental design of the RNAseq screen. Cultured CHRNA3-EGFP^+^ DRG neurons, which acquire mechanotransduction currents upon NGF treatment (top traces), were collected by aspiration into a patch-pipette and subsequently processed for RNA sequencing. **b**, Volcano plot showing the mean fold change of expression (log_2_FC) vs. the log P-Value (N = 3 samples per condition; 20 cells per sample, Student’s T-test). **c**, Comparison of the expression levels (counts per million, CPM) of peptidergic nociceptor subclass markers and **d** mechanically-gated ion channels and PIEZO2 modulators, determined by RNAseq in CHRNA3-EGFP^+^ neurons culture with and without NGF. **e**, Comparison of the mean ± SEM expression levels of TMEM100 normalized to the expression levels of the housekeeping gene GAPDH in the indicated nociceptor subclasses, cultured in the absence and presence of NGF. To enable identification of peptidergic C-fiber nociceptors, non-peptidergic C-fiber nociceptors and Aδ-fiber nociceptors for sample collection, cultures were prepared from Tg(Npy2r-cre)SM19Gsat/Mmucd x B6;129S-Gt(ROSA)26Sortm32(CAG-COP4*H134R/EYFP)Hze/J (Npy2r^Cre^;ChR2-EYFP) mice, in which Aδ-fiber nociceptors express EYFP and were additionally labelled with Alex-Fluor-568 conjugated Isolectin B4 (IB4), which selectively binds to non-peptidergic C-fiber nociceptors. Numbers of samples (20 cells each) per cell type are indicated above the bars and individual values are shown as black dots (Mann-Whitney test: ns, P>0.05, ***, P=0.000082). **f**, Image showing cultured DRG neurons from CHRNA3-EGFP mice transfected with TMEM100-dsRed. **g**, Cartoon depicting the mechano-clamp configuration of the patch clamp technique (left), example traces of mechanically-evoked currents in CHRNA3-EGFP^+^ control cells (middle, top) and in TMEM100-dsRed transfected CHRNA3-EGFP^+^ cells (middle, bottom) as well as bar graph showing the proportion of cells responding to mechanical stimulation. Proportions were compared with Fisher’s exact test. (P=0.023). **h**, Mean ± SEM peak amplitudes of mechanically-evoked currents are shown as a function of membrane displacement for control (white circles) and TMEM100 transfected cells (red squares). Current amplitudes were compared using multiple Mann-Whitney tests (*, P<0.05; **, P<0.01). N-numbers differ from **g**, because some recordings crashed before maximal mechanical stimulation. Source data and statistical information are provided as a Source Data file.

We thus next asked if the up-regulation of TMEM100 is involved in the acquisition of mechanosensitivity in MIAs. To this end we compared mechanotransduction currents from control and TMEM100-overexpressing CHRNA3-EGFP^+^ DRG neurons using an electrophysiological approach known as the mechano-clamp technique ^30^. Here, transmembrane currents are recorded from cultured DRG neurons in the whole-cell configuration of the patch-clamp technique while the cell soma is mechanically stimulated with a fire-polished patch-pipette. Consistent with our previous results ^19^, only a small proportion of un-transfected CHRNA3-EGFP^+^ cells (3/14) responded to mechanical stimulation with small inward currents (Fig. 1f – h). Strikingly, however, ∼61% of the CHRNA3-EGFP^+^ neurons transfected with TMEM100 exhibited robust mechanotransduction currents that were significantly bigger than the small currents occasionally observed in control cells (Fig. 1g and h). When expressed in HEK293 cells, TMEM100 did not produce mechanotransduction currents and did not modulate PIEZO2 mediated currents in these cells (Supplementary Fig. 1), indicating that TMEM100 is neither a channel itself nor a modulator of PIEZO2, but solely un-silences PIEZO2 in the specific cellular context of CHRNA3-EGFP^+^ MIAs.

### Intraarticular CFA injection induces knee joint pain and secondary allodynia in remote skin regions

To corroborate our in-vitro observations, we next examined the role of TMEM100 in the sensitization of MIAs in an *in-vivo* mouse model of Complete Freund’s adjuvant (CFA)-induced knee joint monoarthritis. We chose knee joint inflammation as the experimental model because (i) MIAs were shown to constitute ∼50% of all articular nociceptive afferents ^11,19^, (ii) because the levels of NGF, which induces up-regulation of TMEM100, are significantly increased in synovial fluid in rodent models of inflammatory knee joint pain as well as in patients with osteoarthritis ^31,32^ and (iii) because anti-NGF antibodies alleviate joint pain in patients with osteoarthritis ^33^, suggesting that NGF and possibly MIAs, may play an important role in the generation of knee joint pain.

Consistent with our previous study ^19^, we found that the knee joint is densely innervated by CHRNA3-EGFP^+^ afferents that co-express calcitonin gene-related peptide (CGRP), which mostly terminate in Hoffa’s fat pad (Fig. 2a – c). As previously described ^34^, intraarticular CFA injection caused prominent knee joint inflammation characterized by redness and swelling (Fig. 2d), which was accompanied by severe limping (Supplementary Movies S1 and S2) – indicative of primary hyperalgesia in the knee – and by secondary mechanical and thermal hypersensitivity in skin regions remote from the knee joint. Primary knee joint hyperalgesia was quantified with the Catwalk XT gait analysis system (Fig. 2e), which revealed that mice with an inflamed knee joint put less weight on the affected leg, evidenced by a reduction of the ratio of the foot print area of the ipsi-(left) and contralateral (right) hind paw (before, 1.07 ± 0.03 vs. 3 days post injection (dpi) CFA, 0.57 ± 0.07, N=16, Students paired t-test, P= 2.6*10^−7^), and a reduction of the leg swing speed of the inflamed leg (before, 1.041 ± 0.017 vs. 3 dpi CFA, 0.634 ± 0.044, N=16, Students paired t-test, P= 8*10^−9^ ; Fig. 2f). Secondary mechanical and thermal hypersensitivity was assessed in the ipsilateral hind paw using the von Frey and Hargreaves tests, respectively. These tests showed that CFA-induced monoarthritis, reduces the minimal force of punctate mechanical stimuli – applied with von Frey filaments to the plantar surface of the hind paw – required to evoked a paw withdrawal reflex from 0.87 ± 0.03 g before to 0.15 ± 0.03 g three days post CFA injection (Student’s paired t-test, P=5.18*10^−12^; Fig. 2g). Likewise, the latencies of paw withdrawals evoked by heat stimulation of the hind paw were also significantly reduced (before, 6.03 ± 0.28 s vs. 3 dpi CFA 2.31 ± 0.11 s N=16, Student’s paired t-test, P=1.8*10^−9^, Fig. 2h).

**Fig. 2.**
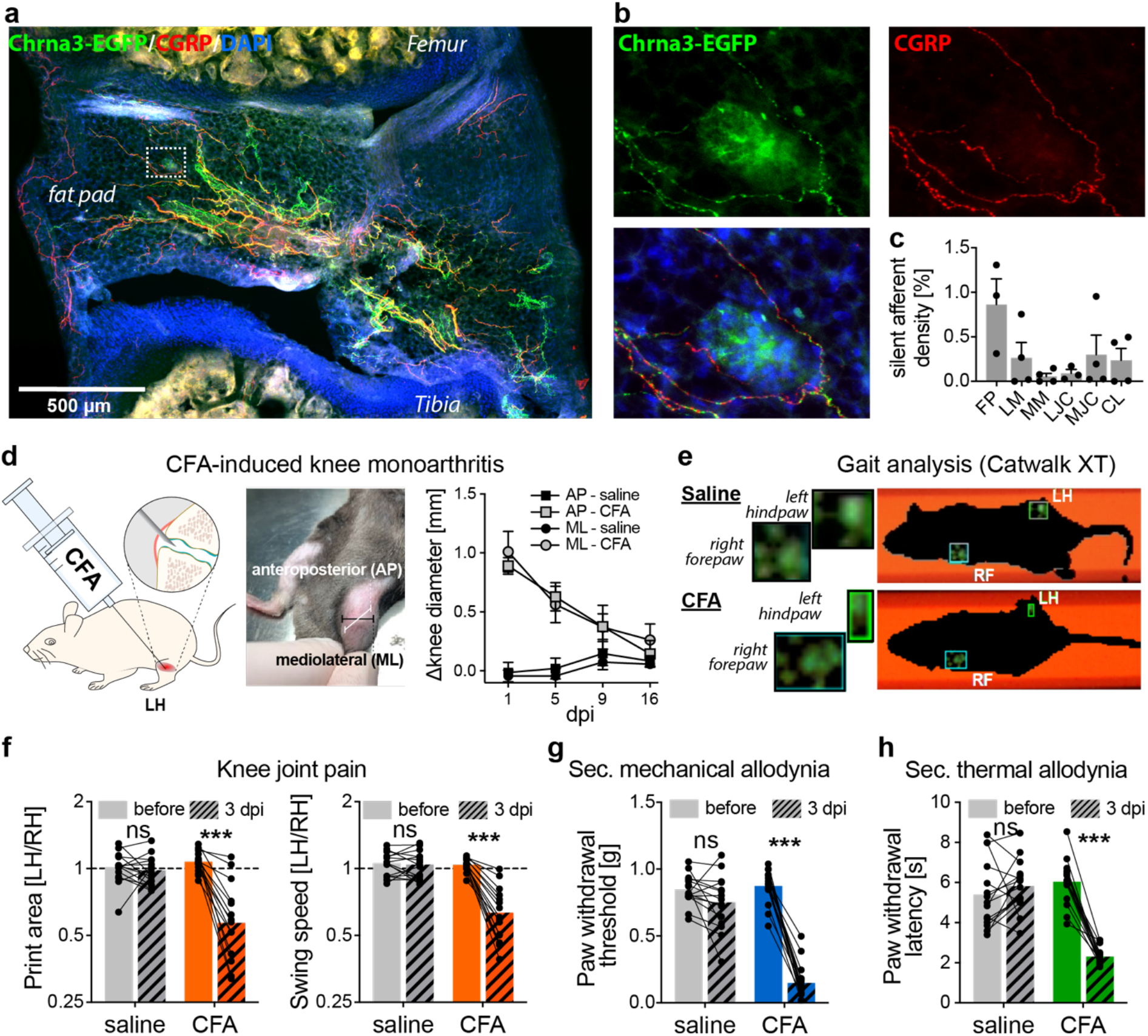
Intraarticular CFA injection induces knee joint pain and secondary allodynia in the hind paw. **a**, Representative image of a knee joint section immunostained for CGRP and EGFP to enhance the endogenous EGFP signal of CHRNA3^+^ afferents. Note, only EGFP^+^ fibers that co-express CGRP are sensory afferents. CGRP^−^/EGFP^+^ fibers are sympathetic efferents (see Prato et al., 2017). **b**, Close-up of the region marked by the white rectangle in (A), emphasizing co-expression of EGFP and CGRP. **c**, Quantification of silent afferent density (EGFP^+^/CGRP^+^ fibers) in anatomically defined regions. FP (Hoffa’s fat pad), LM (lateral meniscus), MM (medial meniscus), LJC (lateral joint capsule), MJC (medial joint capsule), CL (cruciate ligament). Bars represent means ± SEM from three different mice. Individual values from each mouse are shown as black dots. **d**, Cartoon depicting the experimental approach (left), photograph of an inflamed knee (middle) and time course (days post-injection, dpi) of the inflammation-induced knee swelling (right). **e**, Freeze frames of Catwalk XT movies from saline (top) and CFA (bottom) treated mice. Note, CFA-treated mice only put little weight on the left hind paw (small foot print). **f**, Comparison of the foot print area ratio (left/right hindpaw; LH/RH) and the leg swing time ratio (LH/RH) measured before (solid bars) and three day after (3 dpi, hatched bars) saline (grey) and CFA (orange) injection. Paired Student’s t-test (saline N=15, CFA N=16; print area CFA, P = 2.6*10^−7^; swing speed CFA P=8*10^−9^). **g**, Comparison of mechanical paw withdrawal thresholds before (solid bars) and three day after (3 dpi, hatched bars) saline (grey) and CFA (blue) injection. Paired Student’s t-test (saline N=15, CFA N=16; CFA, P = 5.18*10^−12^). **h**, Comparison of thermal paw withdrawal latencies before (solid bars) and three day after (3 dpi, hatched bars) saline (grey) and CFA (blue) injection. Paired Student’s t-test (saline N=15, CFA N=16; CFA, P = 1.8*10^−9^). Source data and statistical information are provided as a Source Data file.

### CFA-induced knee joint inflammation induces mechanosensitivity and potentiates TRPA1 activity in CHRNA3-EGFP^+^ afferents

To enable the examination of CFA-induced transcriptional and functional changes in MIAs, we next back-labelled sensory neurons that give rise to articular afferents by intraarticular injection of the retrograde tracer Fast Blue (FB) (Fig. 3a). Quantification of FB^+^ cells in serial sections of L3 and L4 DRGs, showed that this approach labelled a total of ∼340 DRG neurons (191.5 ± 39.8 cells in L3 DRGs and 150.3 ± 56.7 cells in L4 DRGs, Fig. 3a). IB4-labelling of DRG cultures from FB-injected mice, further showed that 35.1% (108/308 FB^+^ cells) of the FB^+^ cells were CHRNA3-EGFP^+^, 25.3 % (78/308 FB^+^ cells) were small diameter (<30 μm) IB4^−^ peptidergic nociceptors and 26 % (80/308 FB^+^ cells) were large diameter neurons (Fig. 3b) that most likely give rise to group II articular afferents that detect innocuous stimuli. Only a small proportion of the retrogradely labeled neurons were IB4^+^ (13.6%, Fig. 3b), demonstrating that the great majority of nociceptive knee joint afferents are peptidergic (IB4^−^ and CHRNA3-EGFP^+^) and that CHRNA3-EGFP^+^ neurons account for ∼47 % (108/228 FB^+^ nociceptors) of all articular nociceptive afferents. A comparison of the TMEM100 expression levels in small diameter (<30 μm) IB4^−^/FB^+^ and CHRNA3-EGFP^+^/FB^+^ cells from the ipsi- and contralateral L3 and L4 DRGs showed that, similar to *in-vitro* NGF treatment, CFA-induced knee joint inflammation selectively up-regulates TMEM100 in CHRNA3-EGFP^+^ MIAs but not in other C-fiber nociceptors (Fig. 3c).

**Fig. 3.**
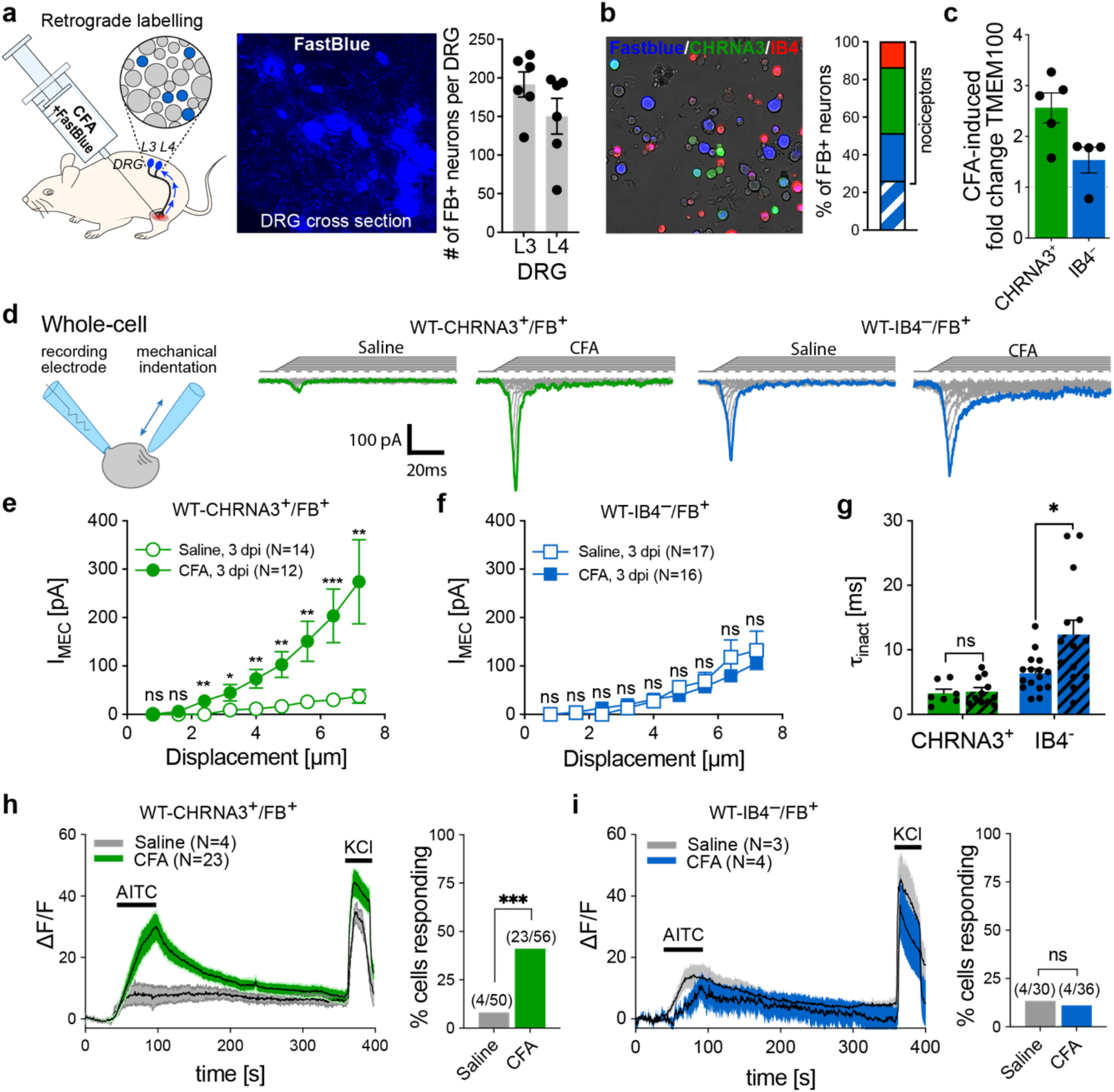
CFA-induced knee joint inflammation induces mechanosensitivity and potentiates TRPA1 activity in CHRNA3-EGFP^+^ afferents. **a**, Cartoon depicting the retrograde labeling approach of knee joint afferents (left), example image of fast blue positive neurons in a DRG cross section (middle) and quantification of the total number of FB^+^ neurons in L3 and L4 DRGs from 6 mice. Bars represent means ± SEM and individual values for each DRG are shown as black dots. **b**, Example image of a primary DRG culture from a CHRNA3-EGFP mouse that had received intraarticular FB (left). The stacked bar graph (right) shows the proportions of IB4^+^ cells (non-peptidergic nociceptors, red), MIAs (green), peptidergic nociceptors (IB4^−^, < 30μm, blue) and group II articular afferents (IB4^−^, >30μm, blue hatched). **c**, Quantification of CFA-induced changes in TMEM100 expression in retrogradely labelled MIAs and peptidergic nociceptors. Expression levels in ipsi- and contralateral DRGs were determined by qPCR and compared using the ΔΔCt method. **d**, Cartoon depicting the mechano-clamp configuration of the patch clamp technique (left) and example traces of mechanically-evoked currents in the indicated cell types and conditions (right). **e** and **f** Peak amplitudes of mechanically-evoked currents are shown as a function of membrane displacement for **e** MIAs and **f** peptidergic nociceptors from saline (open symbols) and CFA-treated (solid symbols) mice. Current amplitudes were compared using Mann-Whitney test (*, P<0.05; **, P<0.01; ***, P<0.001; ns, P>0.05). **g**, Comparison of the inactivation time constants of the mechanically evoked currents determined with single exponential fit. Bars represent means ± SEM and individual values are shown as black dots. **h** and **i** Time course of changes in the intracellular Ca^2+^ concentration determined with Calbryte-590 Ca^2+^ imaging (left) of **h** retrogradely labelled MIAs and **i** peptidergic nociceptors from saline and CFA treated mice, responding to 10 μM AITC and 100 mM KCl and bar graphs (right) showing the proportion of cells that responded to AITC. N-numbers are provided in the graph legends. Proportions were compared with Fishers exact test. ns, not significant; ***, P=0.000102. Source data and statistical information are provided as a Source Data file.

We next asked if mechanosensitivity of FB-labelled DRG neurons is altered in CFA-induced monoarthritis. In accordance with our previous results ^19^, CHRNA3-EGFP^+^/FB^+^ neurons from saline injected animals did not show currents in response to mechanical stimulation of the plasma membrane (Fig. 3d, e). Following intraarticular CFA-injection (3 dpi), however, FB-labelled CHRNA3-EGFP^+^ neurons exhibited robust mechanotransduction currents that were significantly larger than the small inward currents occasionally observed in control animals (Fig. 3d, e). Interestingly, the amplitudes of the mechanotransduction currents of small-diameter IB4^−^ nociceptors were not altered in CFA-treated mice (Fig. 3f), but we observed a small, yet significant, increase in the inactivation time constants of these currents (Fig. 3g). Since TMEM100 was previously shown to potentiate the activity of TRPA1 ^28^, we also examined the responsiveness of FB-labelled neurons to the TRPA1 agonist allylisothiocyanate (AITC) using Calbryte-590 Ca^2+^-imaging. Strikingly, both the amplitude of the AITC-evoked calcium transients as well as the proportion of cells that respond to AITC, were significantly increased amongst MIAs from CFA-treated mice (Fig. 3h), but were completely unaffected in small diameter IB4^−^ nociceptors (Fig. 3i).

Taken together, our data showed that intraarticular CFA-injection causes knee joint pain and secondary hyperalgesia in the ipsilateral hind paw, which is accompanied by an upregulation of TMEM100, the potentiation of TRPA1 activity and, most importantly, the acquisition of mechanosensitivity in CHRNA3-EGFP^+^ neurons.

### TMEM100 knock-out mice develop normal inflammatory knee joint pain but no long-lasting secondary mechanical allodynia

We next asked if the pain phenotype of CFA-induced knee monoarthritis (Fig. 2f – h) is causally linked to the sensitization of CHRNA3-EGFP^+^ nociceptors and if this sensitization is induced by the upregulation of TMEM100. To this end we generated conditional TMEM100 knock-out mice, hereafter referred to as TMEM100KO mice, by crossing mice that carry a conditional allele for TMEM100 ^35^ with SNS-Cre mice, in which Cre-recombinase expression is driven by the voltage-gated sodium channel Na_v_1.8 promoter ^36^ and is thus expressed in all nociceptors including CHRNA3-EGFP^+^ neurons ^37^. We first compared primary knee joint pain, assessed by gait analysis, in male wildtype (WT) mice that received intraarticular saline injections with male WT and TMEM100KO mice that received CFA injections, over a period of 21 days. WT mice that received CFA, exhibited significantly altered gait, indicative of knee joint pain, during the first seven days post CFA injection compared to saline treated animals. Surprisingly, CFA-treated TMEM100KO mice also developed knee joint pain and showed altered gait (Supplementary movie S3), which significantly differed from saline-treated WT mice (Supplementary movie S1) and was indistinguishable from gait observed in CFA-treated WT mice (Supplementary movie S2; Fig. 4a). Strikingly, however, mechanical allodynia in the ipsilateral hind paw was significantly attenuated in TMEM100KO mice. Thus, the mechanical paw withdrawal thresholds were only transiently reduced from day 3 until day 5 and returned to baseline values by day 7 in TMEM100KO mice, while WT mice exhibited long-lasting secondary mechanical allodynia, which persisted until the end of the examination period (21 dpi, Fig. 4b). Secondary thermal hypersensitivity was not altered in TMEM100KO mice (Fig. 4c).

**Fig. 4.**
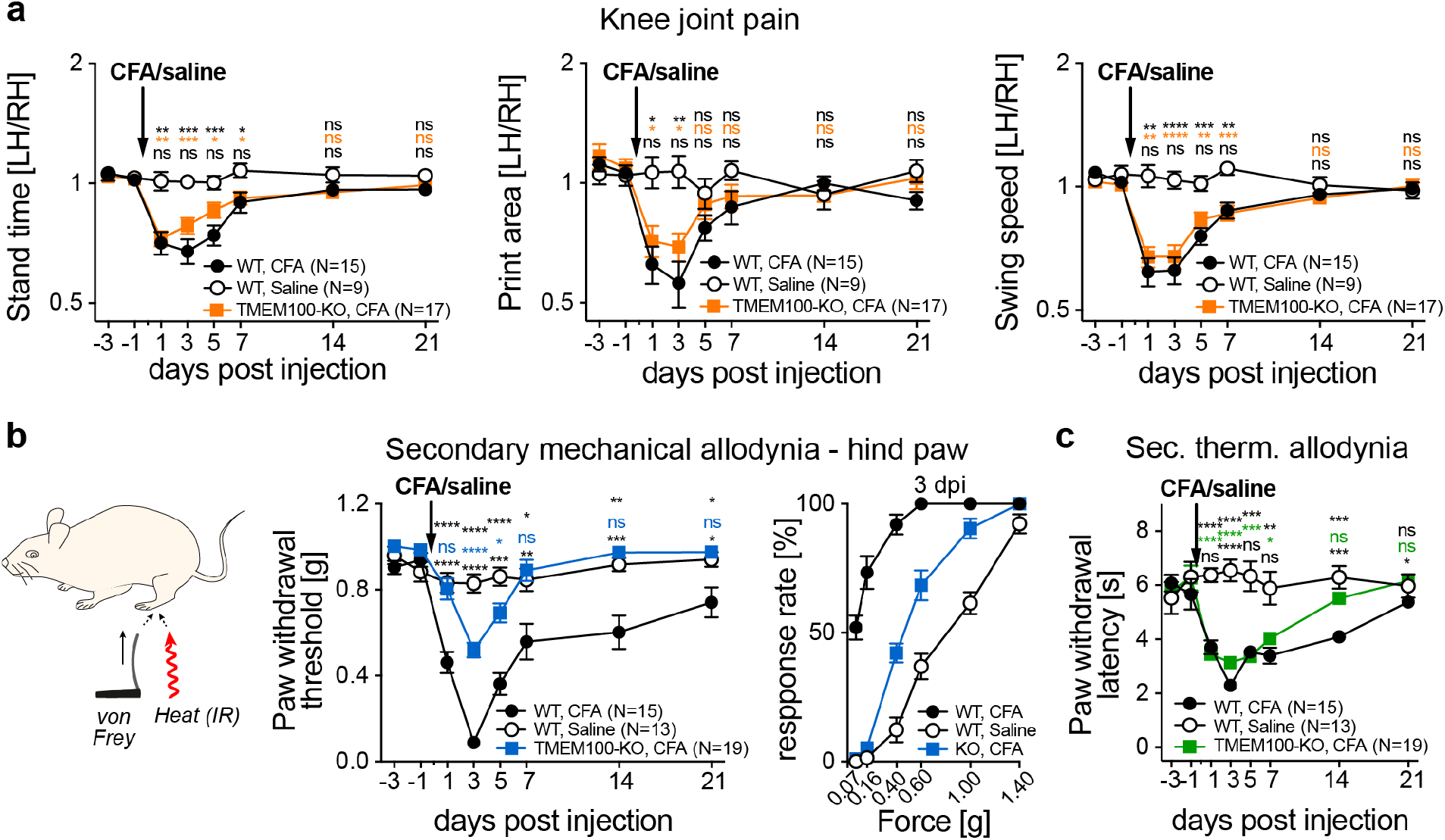
TMEM100 knock-out mice develop normal inflammatory knee joint pain but no long-lasting secondary mechanical allodynia. **a**, Comparison of the time courses of changes in stand time (left), foot print area (middle) and leg swing speed (right) of saline injected WT mice (white circles), CFA injected WT mice (black circles) and CFA injected TMEM100KO mice (orange squares). **b**, Cartoon depicting the experimental approach for measuring secondary mechanical and thermal hypersensitivity in the ipsilateral hindpaw (left), time courses of changes in mechanical paw withdrawal thresholds of saline injected WT mice (white circles), CFA injected WT mice (black circles) and CFA injected TMEM100KO mice (blue squares) (middle) and responsiveness of WT mice (white circles), CFA injected WT mice (black circles) and CFA injected TMEM100KO mice (blue squares) three days post saline/CFA injection (3 dpi) to all tested von Frey filaments (right). Response rates were compared using 2-way ANOVA. P-values of multiple comparisons are as follows: WT-saline vs. WT-CFA, P_0.07g_ = 7.5E-08, P_0.16g_ = 2.8E-08, P_0.4g_ = 8.7E-12, P_0.6g_ = 7.5E-08, P_1g_ = 2.6E-06, P_1.4g_ = 1.2E-01; WT-saline vs. TMEM100KO-CFA, P_0.07g_ = 5.9E-01, P_0.16g_ = 3.3E-01, P_0.4g_ = 1.5E-04, P_0.6g_ = 7.8E-04, P_1g_ = 6.8E-05, P_1.4g_ = 1.2E-01; WT-CFA vs. TMEM100KO-CFA, P_0.07g_ = 5.2E-08, P_0.16g_ = 3.6E-08, P_0.4g_ = 3.9E-10, P_0.6g_ = 9.9E-05, P_1g_ = 6.1E-02, P_1.4g_ = not determined). **c**, Time courses of changes in thermal paw withdrawal latencies of saline injected WT mice (white circles), CFA injected WT mice (black circles) and CFA injected TMEM100KO mice (green squares). **a** – **c** Symbols represent means ± SEM. Unless otherwise stated, ratios at different time points were compared using mixed model ANOVA and P-values of multiple comparisons are indicated above the white circles. Top, WT-saline vs. WT-CFA; middle, WT-saline vs. TMEM100KO-CFA; bottom, WT-CFA vs. TMEM100KO-CFA (*, P < 0.05; **, P < 0.01; ***, P < 0.001; ns, P > 0.05, N-numbers are provided in the graph legends). Source data and statistical information are provided as a Source Data file.

Since an increasing body of literature demonstrates sex differences with regards to pain sensitivity, we reproduced the behavioral experiments using female TMEM100KO mice. Interestingly, female mice exhibited the exact same pain phenotype as male mice with respect to primary and secondary hypersensitivity (Supplementary Fig. 2) indicating a sex-independent role of TMEM100 in inflammatory pain.

### CFA-induced knee joint inflammation fails to sensitize CHRNA3-EGFP^+^ neurons to mechanical stimuli in TMEM100 knock-out mice

We next examined the role of TMEM100 in the acquisition of mechanosensitivity and the potentiation of TRPA1 activity in CHRNA3-EGFP^+^ knee joint afferents in CFA-induced monoarthritis. Patch-clamp recordings showed that articular MIAs, (FB-labelled CHRNA3-EGFP^+^ neurons) from TMEM100KO mice did not acquire mechanosensitivity during CFA-induced inflammation (Fig. 5a, b) and that mechanosensitivity of FB^+^/IB^−^ neurons was also not altered (Fig. 5a – d). Moreover, in accordance with the previously described interaction between TMEM100 and TRPA1 ^28^, AITC-induced TRPA1-mediated Ca^2+^-signals were not potentiated in CHRNA3-EGFP^+^ knee joint afferents from TMEM100KO mice (Fig. 5e), nor were the TRPA1 responses of small diameter IB4^−^ nociceptors altered (Fig. 5f).

**Fig. 5.**
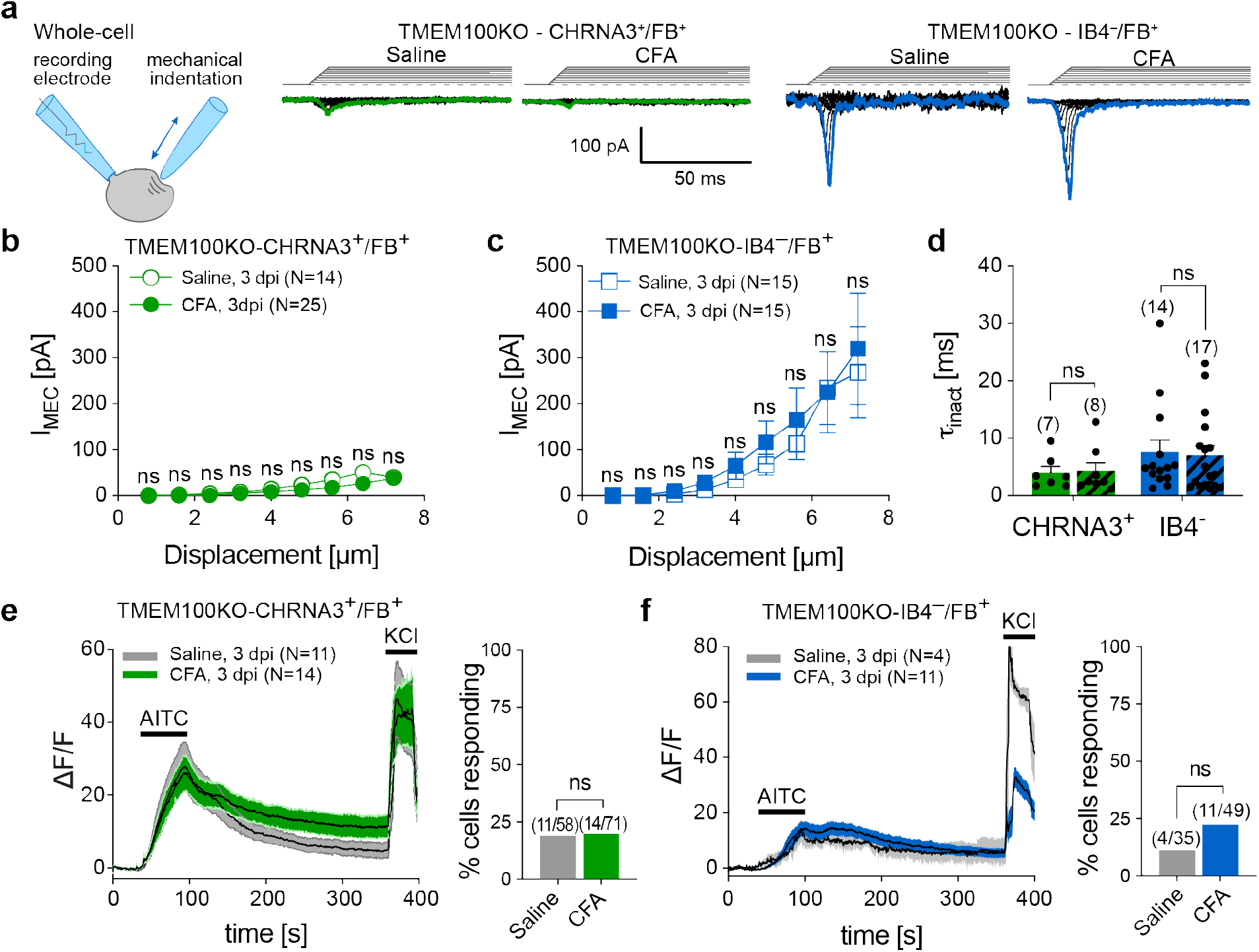
CFA-induced knee joint inflammation fails to sensitize CHRNA3-EGFP^+^ neurons to mechanical stimuli in TMEM100 knock-out mice. **a**, Cartoon depicting the mechano-clamp configuration of the patch clamp technique (left) and example traces of mechanically-evoked currents in the indicated cell types and conditions (right). Note, in contrast to WT mice, CHRNA3-EGFP^+^ neurons from TMEM100KO mice do not acquire mechanosensitivity during CFA-induced inflammation. **b** and **c** Peak amplitudes of mechanically-evoked currents are shown as a function of membrane displacement for **b** MIAs and **c** peptidergic nociceptors from saline (open symbols) and CFA-treated (solid symbols) mice. Current amplitudes were compared using Mann-Whitney test (ns, P>0.05). **d**, Comparison of the inactivation time constants of the mechanically evoked currents determined with single exponential fit. Bars represent means ± SEM and individual values are shown as black dots. **e** and **f** Time course of changes in the intracellular Ca^2+^ concentration determined with Calbryte-590 Ca^2+^ imaging (left) of retrogradely labelled **e** MIAs and **f** peptidergic nociceptors from saline (grey) and CFA (green) treated mice, challenged with 10 μM AITC and 100 mM KCl and bar graphs (right) showing the proportion of cells that responded to AITC. N-numbers are provided in the graph legends. Proportions were compared with Fishers exact test (ns, not significant). Black lines represent means and grey and green shaded areas represent SEM. Source data and statistical information are provided as a Source Data file.

Hence, our data indicates that the un-silencing of silent nociceptors – i.e. the acquisition of mechanosensitivity and the potentiation of TRPA1 – strictly depends on the up-regulation of TMEM100 expression. Considering that secondary mechanical allodynia in the hind paw, but not knee joint pain, was greatly attenuated in TMEM100KO mice during CFA-induced knee inflammation (Fig. 4), together with the observation that the lack of TMEM100 solely altered mechanosensitivity and TRPA1 function in CHRNA3-EGFP^+^ afferents, but not in IB4^−^ peptidergic knee joint afferents (Fig. 3 and Fig. 5), we hypothesized that the un-silencing of silent nociceptors triggers the development of secondary mechanical allodynia.

### Sensitization of cutaneous C-fiber nociceptors contributes to secondary mechanical allodynia

It is well established that secondary mechanical allodynia results from central sensitization – i.e. a strengthening of synaptic transmission in pain processing circuits in the spinal cord ^38,39^. Considering the remarkable reduction of paw withdrawal thresholds in CFA-induced knee joint monoarthritis (Fig. 4b), however, we asked if sensitization of cutaneous nociceptors also contributes to secondary mechanical allodynia. To examine peripheral sensitization, we directly measured the mechanosensitivity of C-fiber and Aδ-fiber nociceptors in the tibial nerve, which innervates the plantar surface of the hind paw, by recording mechanically evoked action potentials from single nerve fibers in an ex-vivo skin-nerve preparation from mice that had received intraarticular CFA (Fig. 6a, b and Supplementary Fig. 3). These recordings revealed, that 40% of the cutaneous C-fiber nociceptors from mice with CFA-induced knee monoarthritis, are activated by forces of 0.16 g and below and virtually all C-fibers (90%) responded to von Frey filaments of 0.4 g and below (Fig. 6a – d). By contrast, not a single C-fiber nociceptor from saline-treated control mice responded to von Frey stimuli ≤ 0.16g and only ∼17% were activated by the 0.4 g filament (Fig. 6a – d). Moreover, cutaneous C-fibers from CFA-treated mice also fired significantly more action potentials in response to suprathreshold mechanical ramp-and-hold stimuli applied with a piezoelectric micromanipulator (Fig. 6e). Aδ-fiber nociceptors also showed a trend towards being more sensitive, but this difference was less pronounced than in C-fibers (Supplementary Fig. 3b). Likewise, the firing rate was slightly higher in Aδ-fiber nociceptors from CFA-treated mice, but only significant for the largest tested suprathreshold stimulus (Supplementary Fig. 3c). Strikingly, the mechanical activation thresholds and the firing rates of both C-fiber nociceptors and Aδ-fiber nociceptors from CFA-treated TMEM100KO mice were indistinguishable from those of saline-treated WT control mice (Fig. 6d, e and Supplementary Fig. 3b and c). Taken together, these results show that CFA-induced knee joint inflammation shifts the mechanosensitivity of individual cutaneous C-fiber nociceptors towards innocuous mechanical stimuli, which correlates with the leftward shift of the paw withdrawal thresholds in the same mice. Hence, our data suggests that in addition to central sensitization, sensitization of cutaneous C-fiber and Aδ-fiber nociceptors might also significantly contribute to secondary mechanical allodynia in the hind paw induced by knee joint inflammation.

**Fig. 6.**
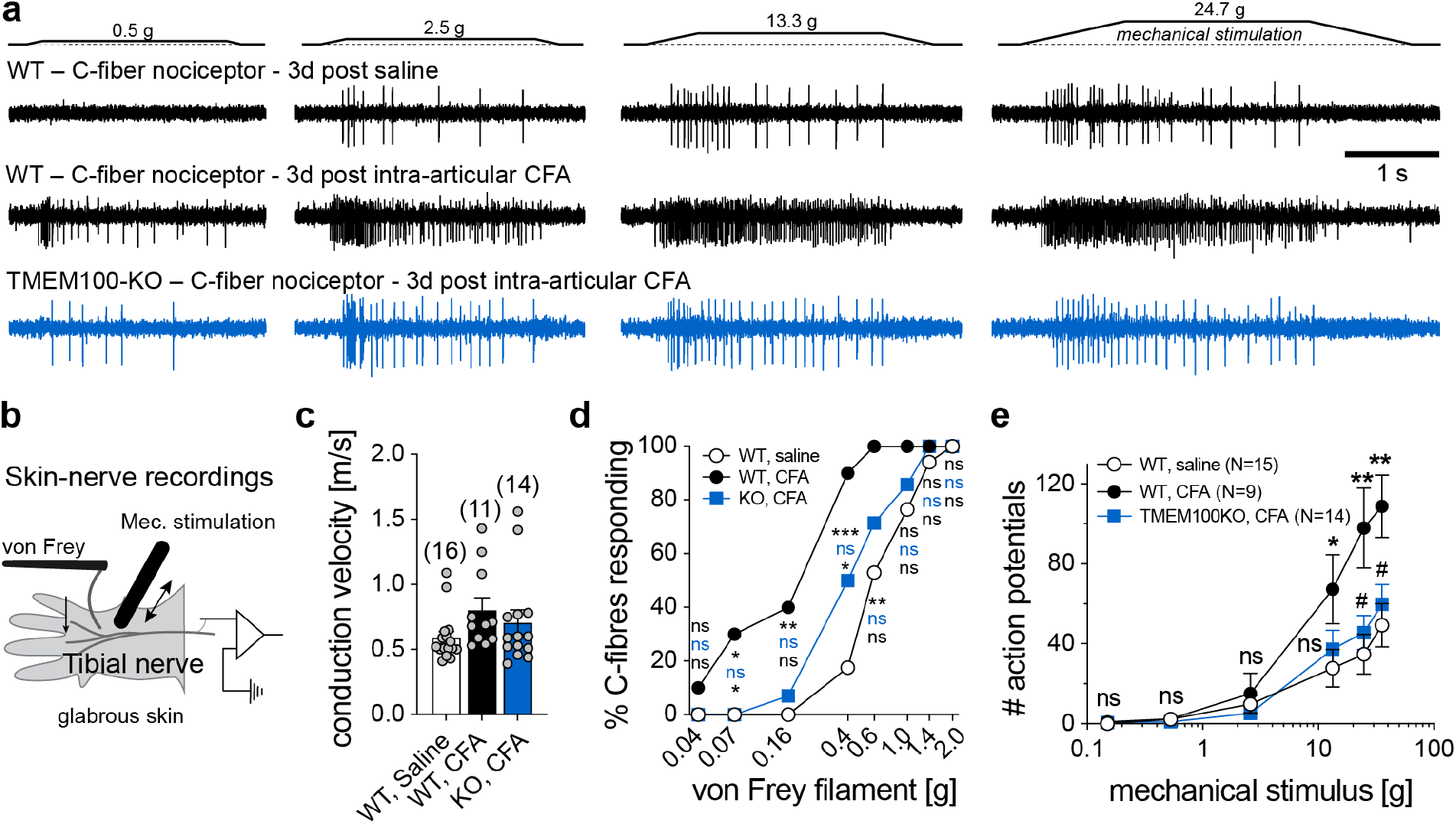
Sensitization of cutaneous C-fiber nociceptors contributes to secondary mechanical allodynia. **a**, Example traces of mechanically-evoked action potentials recorded from single nerve fibers from the tibial nerve of WT mice 3 days post intraarticular saline (top), WT mice 3 dpi CFA (middle) and TMEM100KO mice 3 dpi CFA (bottom). **b**, Cartoon depicting the experimental approach. **c**, Scatter dot plots of the conduction velocities [m/s] of the examined afferent fibers from the indicated mice. Bars represent means ± SEM. **d**, Comparison of the proportions of C-fiber nociceptors that respond to mechanical stimulation with the indicated von Frey filaments. The proportions were compared pairwise using the Chi-square test. P-values are provided next to the symbols in the graph and refer to WT-saline vs. WT-CFA (top, black), WT-saline vs. TMEM100KO-CFA (middle, blue) and WT-CFA vs. TMEM100KO (bottom, black). N-numbers are the same as in (C). **e**, Comparison of the firing rates evoked by a series of ramp-and-hold stimuli with increasing amplitudes that exerted the indicated force to the receptive fields. Symbols represent the mean ± SEM numbers of action potentials, which were compared using multiple Mann-Whitney tests. (WT-saline vs. WT-CFA: *, P<0.05; **, P<0.01; WT-CFA vs. TMEM100KO-CFA: #, P<0.05). Source data and statistical information are provided as a Source Data file.

### Overexpression of TMEM100 in articular afferents induces secondary allodynia in the hind paw but no knee joint pain

While the spinal and supraspinal mechanisms of central sensitization that underlie secondary hyperalgesia are well understood, only little is known about the initial signals that trigger the development of secondary allodynia ^38,40^. Considering that inhibition of un-silencing of MIAs by knocking out TMEM100 prevents the development of long-lasting secondary allodynia, our data suggests that MIAs might be essential for the induction of secondary allodynia. To test this hypothesis, we next un-silenced articular MIAs without inducing an inflammation. To this end, we selectively overexpressed TMEM100 in knee joint afferents by intra-articular injection of an AAV-PHP.S-TMEM100-Ires-dsRed virus (30 μl, 1.5*10^11^ vg; Fig. 7a). Four days after intraarticular AAV-PHP.S-TMEM100-Ires-dsRed administration, we observed prominent dsRed fluorescence in a total of 339 ± 7 neurons in ipsilateral L3 and L4 DRG (Fig. 7a). Most importantly, we observed numerous dsRed expressing nerve fibers in the saphenous nerve proximal to the knee (Fig. 7b), which includes the medial articular nerve that supplies the knee joint and in which silent nociceptors had first been described ^11^, but hardly any dsRed^+^ fibers in the tibial nerve distal to the knee, which contains cutaneous afferents that supply the plantar surface of the hind paw (Fig. 7b). Hence, intraarticularly administered AAV-PHP.S-TMEM100-Ires-dsRed causes selective overexpression of TMEM100 in knee joint afferents, but not in afferents that supply the plantar surface of the hind paw. Interestingly, TMEM100-overexpressing mice exhibited normal gait, indicating that un-silencing of knee joint MIAs does not trigger knee joint pain (Fig. 7c). Strikingly, however, these mice developed profound mechanical hypersensitivity in the ipsilateral hind paw five days post AAV injection, which persisted until the end of the observation period (21 dpi; Fig. 7d). Thus, the mechanical paw withdrawal thresholds decreased from 0.92 ± 0.066 g (−1 dpi) to 0.373 ± 0.035 g (14 dpi). Mice that received a control virus without TMEM100 (AAV-PHP.S-dsRed) did not show any signs of mechanical hypersensitivity. In accordance with the behavioral outcome of TMEM100 over-expression in articular afferents, single-unit action potential recordings from a tibial nerve-glabrous skin preparation showed that cutaneous C-fiber nociceptors fire about twice as many action potentials in response to a given stimulus and have significantly reduced mechanical activation thresholds in mice that overexpress TMEM100 in articular afferents compared to mice that had received a control virus (Fig. 7e – g). Similar to mice that received intraarticular CFA, the mechanical activation thresholds of cutaneous Aδ-fiber nociceptors were slightly reduced and the action potential firing rate in response to suprathreshold stimuli was significantly increased (Supplementary Fig. 4).

**Fig. 7.**
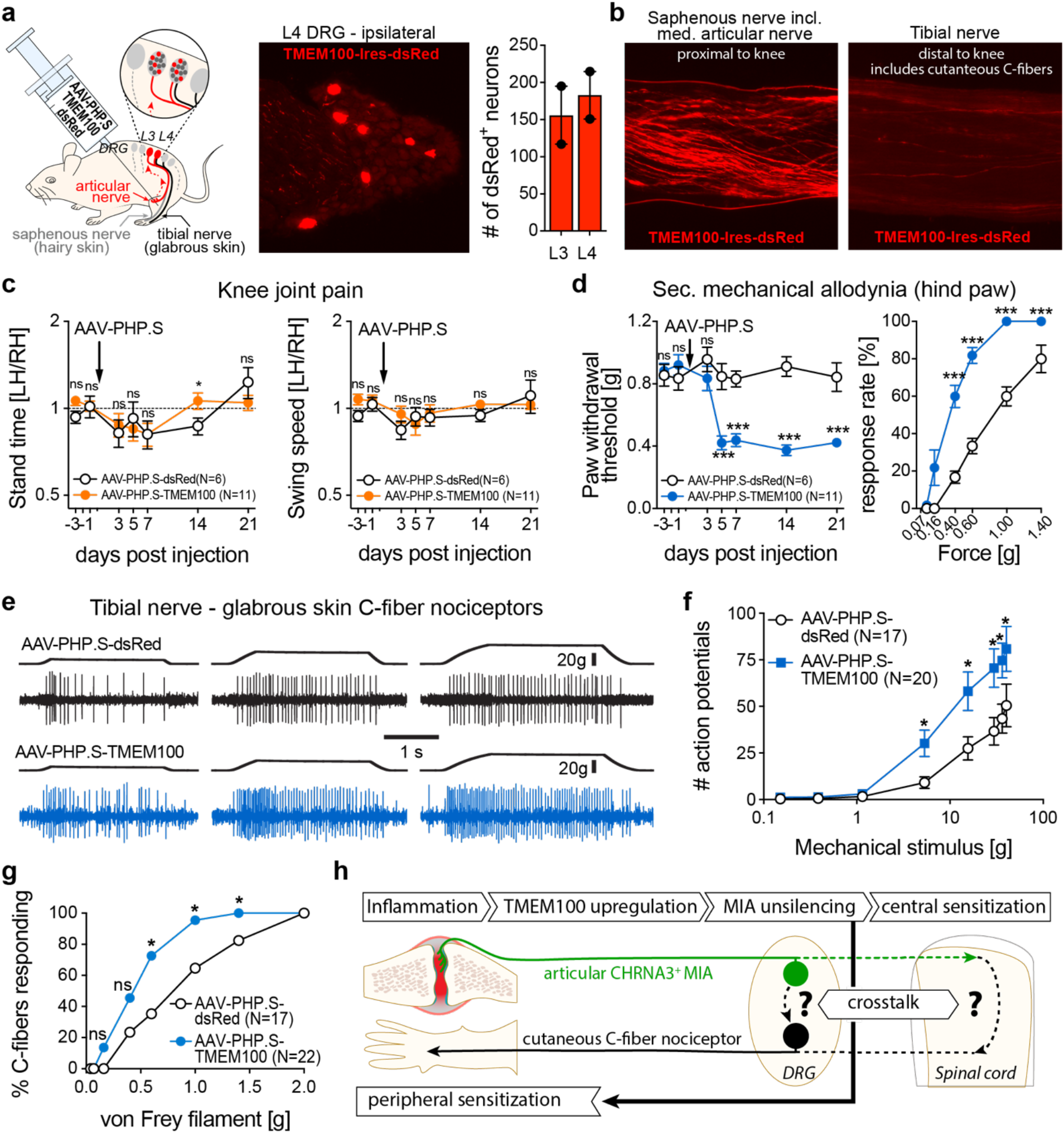
Overexpression of TMEM100 in articular afferents induces secondary allodynia in the hind paw but no knee joint pain. **a**, Cartoon depicting the experimental approach (left), example image of a DRG cross section from a mouse that had received intraarticular AAV-PHP.S-TMEM100-Ires-dsRed (middle) and quantification of the total number of dsRed^+^ neurons (right). Bars represent means ± SEM and values from individual DRGs are shown as black dots. **b**, dsRed fluorescence in the tibial nerve distal to the knee (left) and the saphenous (right) nerve proximal to the knee which contains the medial articular nerve. **c**, comparison of the time courses of changes in stand time (left) and leg swing speed (right) of WT mice that intraarticularly received AAV-PHP.S-dsRed control virus (white circles) and AAV-PHP-S-TMEM100-Ires-dsRed (orange circles). Ratios at different time points were compared using multiple Mann-Whitney tests. (ns, P>0.05; *, P<0.05; N-numbers are provided in the graph legend). **d**, Time courses of changes in mechanical paw withdrawal thresholds of AAV-PHP.S-dsRed (white circles) and AAV-PHP-S-TMEM100-Ires-dsRed (blue circles) injected WT mice (left) as well as comparison of the response rates at 14 dpi – i.e. % paw withdrawals in response to five successive stimulations with the indicated von Frey filaments (right). Mean paw withdrawal thresholds at different time points and response rates to different von Frey filaments were compared using Mann-Whitney test (***, P<0.001; number of tested animals is the same in both panels and is indicated in the graph legend). **e**, Example traces of mechanically-evoked action potentials recorded from cutaneous C-fiber nociceptors in the tibial nerve from control mice (top, AAV-PHP.S-dsRed) and from mice that overexpress TMEM100 in articular afferents (bottom, AAV-PHP.S-TMEM100-Ires-dsRed). **f**, comparison of the firing rates evoked by ramp- and-hold stimuli that exerted the indicated force to the receptive fields. Symbols represent means ± SEM numbers of action potentials, which were compared using multiple Mann-Whitney tests (*, P<0.05). **g**, comparison of the proportions of C-fiber nociceptors that respond to mechanical stimulation with the indicated von Frey filaments. The proportions were compared pairwise using the Chi-square test (ns, not significant; *, P<0.05). **h**, cartoon depicting the proposed mechanism underlying the induction of secondary hypersensitivity. Source data and statistical information are provided as a Source Data file.

In summary, our data shows that TMEM100 overexpression-induced un-silencing of mechanically insensitive articular afferents is sufficient to trigger mechanical hypersensitivity in remote skin regions (Fig. 7d). Together with the observation that TMEM100 is specifically up-regulated in MIAs during CFA-induced inflammation and that knock-out of TMEM100 exclusively abolishes long-lasting secondary allodynia, these results suggest that sensory input from unsilenced MIAs is the main trigger for the induction of secondary allodynia (Fig. 7h). While we have not examined the well-established contribution of central sensitization, our data reveals a previously unrecognized contribution of peripheral sensitization to secondary allodynia.

## Discussion

The peculiar properties of MIAs have fueled speculations about a prominent contribution to inflammatory pain ever since they had first been described more than thirty years ago ^11,17,41^, but hitherto, neither the molecular mechanism underlying their un-silencing, nor their exact role in pain signaling have been deciphered.

We have previously shown that *in-vitro* NGF-induced un-silencing of MIAs requires *de-novo* gene transcription and that mechanosensitivity in MIAs is mediated by the mechanically-gated ion channel PIEZO2, the expression of which is, however, not changed during un-silencing (Fig. 1d) ^19^. Here we show that TMEM100 is specifically up-regulated in CHRNA3-EGFP^+^ MIAs during inflammation (Fig. 1d, e and Fig. 3c) and demonstrate that over-expression of TMEM100 is sufficient to un-silence MIAs *in-vitro* (Fig. 1f – h), whereas knock-out of TMEM100 prevents un-silencing in a mouse model of CFA-induced monoarthritis (Fig. 5b, d). Considering that CHRNA3-EGFP^+^ MIAs express high levels of PIEZO2 even in conditions in which they are not mechanosensitive ^19^ (Fig. 1d) and that PIEZO2 does not require the presence of an auxiliary subunit for normal function, we hypothesize that PIEZO2 is somehow kept ‘silent’ in these neurons under normal conditions and is primed by the up-regulation of TMEM100. CHRNA3-EGFP^+^ MIAs express four known PIEZO2 inhibitors, namely MTMR2 ^22^, SERCA2 (Atp2a2) ^26^, Annexin A6 (Anxa6) ^25^ and Nedd4-2 ^27^, but since neither of them is down-regulated by NGF (Fig. 1sd) and all four are ubiquitously expressed in sensory neurons ^2^, it seems highly unlikely that any of them is involved in keeping PIEZO2 silent in MIAs. Hence, while our study identified TMEM100 as a key protein that is sufficient and required for the un-silencing of MIAs, the mechanism by which PIEZO2 is kept inactive in CHRNA3-EGFP^+^ MIAs remains elusive.

TMEM100 had originally attracted our attention because it was previously shown to enhance TRPA1 activity in a TRPV1-dependent manner ^28^ and because there is ample evidence supporting a role of TRPA1 in fine-tuning mechanosensitivity of sensory afferents. Thus, it was shown that pharmacological blockade as well as knock-out of TRPA1 partially inhibit mechanotransduction currents in cultured DRG neurons and reduce the firing rate of cutaneous C-fiber nociceptors ^42–45^. Since TRPA1, to the best of our knowledge, is not activated by mechanical indentation of the cell membrane but is directly gated by intracellular calcium ^46^, a possible mechanistic explanation for the role of TRPA1 is that it increases mechanosensitivity of sensory neurons by amplifying PIEZO2-mediated Ca^2+^ influx. Consistent with the findings of Weng and colleagues ^28^, we found that CFA-induced knee joint inflammation enhances TRPA1 activity in a TMEM100-dependent manner (Fig. 3h and Fig. 5e). Most importantly, our data substantiate and extend the findings of Weng et al. (2015), by showing that TMEM100 is exclusively up-regulated in CHRNA3-EGFP^+^ neurons – at least in CFA-induced knee monoarthritis and after in-vitro NGF treatment (Fig. 1d, e and Fig. 3c) – and accordingly only enhances TRPA1 activity in MIAs but not in other nociceptors (Fig. 3h, i and Fig. 5e, f).

Hence our data suggests that TMEM100 is a pleiotropic protein that on the one hand induces mechanosensitivity in MIAs by priming PIEZO2 via a yet unknown mechanism and on the other hand amplifies PIEZO2-dependent mechanosensitivity by releasing TRPA1 from inhibition by TRPV1.

Primary hyperalgesia has an adaptive role as it produces behaviors that promote healing and recovery. It results from peripheral sensitization – i.e. direct sensitization of nerve endings in the inflamed or injured tissue – and usually resolves concurrently with the initial cause of pain. Secondary allodynia and hyperalgesia, on the other hand, are considered maladaptive and result from central sensitization – i.e. a strengthening of synaptic transmission in pain processing circuits in the spinal cord ^38–40^. Moreover, secondary allodynia is thought to increases the likelihood of developing chronic pain as, for example, the area and intensity of secondary allodynia immediately after surgery correlates with the incidence of persistent postsurgical pain ^47^.

While the spinal mechanisms of central sensitization have been studied extensively in various rodent pain models, including experimentally induced arthritis ^48–50^, the peripheral inputs that initially trigger the development of central sensitization and hence secondary allodynia are still unknown. Here we show that blocking the un-silencing of articular MIAs by knocking out TMEM100 prevents the development of long-lasting secondary allodynia in remote skin regions, but does not alter pain at the actual site of CFA-induced inflammation (Fig. 4). Moreover, TMEM100 overexpression-induced un-silencing of knee joint MIAs in the absence of inflammation or injury, induces mechanical hypersensitivity in the paw but not pain hypersensitivity in the knee joint (Fig. 7c, d). Finally, our skin-nerve recordings demonstrate that secondary allodynia, in addition to the previously described central sensitization, is partly driven by peripheral sensitization of cutaneous C-fiber nociceptors (Fig. 6 and Fig. 7e – g). Regarding the AAV-mediated TMEM100 over-expression experiments, one should note that intra-articularly administered AAV-PHP.S infects all subclasses of sensory afferents in the knee joint and thus the observed secondary allodynia could theoretically also result from TMEM100 over-expression in articular afferents other than CHRNA3-EGFP^+^ MIAs. Although we cannot completely rule out this possibility, a contribution of other nociceptor subtypes to the induction of secondary allodynia seems unlikely, because if ectopic TMEM100 overexpression had also increased the activity other nociceptors in the knee joint, then we should have observed knee joint pain – i.e. changes in gait – in addition to secondary allodynia, which was, however, not the case (Fig. 7c). Moreover, the observation that TMEM100 expression is exclusively upregulated in MIAs in CFA-induced monoarthritis (Fig. 3c) and that mice lacking TMEM100 only show functional deficits in MIAs but not in other articular nociceptors (Fig.3 and 5) and selectively lose secondary allodynia but not knee joint pain, strongly support a specific role of MIAs in inducing secondary allodynia. It should further be noted, that we observed off-target expression of TMEM100 in a few fibers in the tibial nerve distal to the knee, which supplies the plantar surface of the hind paw (Fig. 7b). Cutaneous C-fiber nociceptors that detect von Frey stimuli express MRGPRD ^7^ and require PIEZO2 and TRPA1 for normal mechanosensitivity ^43,52^, but do not express TRPV1 ^2,7^. Since TMEM100, however, only sensitizes PIEZO2 in the specific cellular environment of CHRNA3-EGFP^+^ MIAs (Fig. 1f – h and Supplementary Fig. 1) and enhances TRPA1 in a TRPV1-dependent manner – in fact it inhibits TRPA1 in the absence of TRPV1 ^28^ – the off-target expression of TMEM100 in a few cutaneous nerve fibers most likely does not contribute to the observed secondary hyperalgesia in the hind paw.

We have not explicitly tested if un-silencing of MIAs also triggers central sensitization, which is thought to be the major cause of secondary hypersensitivity. Yet, considering that the AAV-PHP.S-TMEM100 induced reduction of paw withdrawal thresholds (Fig. 7d) was larger than the reduction of the mechanical activation thresholds of individual cutaneous C- and Aδ-fiber nociceptors (Fig. 7g and Fig 4c), it seems highly likely that central processing of nociceptive input was also altered in these experiments such that subliminal sensory inputs evoked pain.

Hence, our data support a mechanistic model of secondary allodynia in which inflammation-induced upregulation of TMEM100 un-silences MIAs, which subsequently triggers central sensitization and – via a yet unknown central mechanism – peripheral sensitization of cutaneous nociceptors, which eventually leads to secondary mechanical allodynia in skin regions remote from the site of inflammation (Fig. 7h). By demonstrating that primary and secondary pain hypersensitivity are triggered by separate subclasses of primary sensory afferents and considering that MIAs constitute almost fifty percent of all nociceptors in viscera and deep somatic tissues, our study provides an invaluable framework for future studies that aim at deciphering the contribution of different afferent subtypes to other clinically relevant forms of pain and to develop new strategies for preventing the transition from acute to chronic of pain after injury, inflammation or surgical interventions.

## Methods

### Animals

All experiments were conducted in accordance with the European Communities Council Directive (EU and institutional guidelines) including the ethical guidelines of ‘Protection of Animals Act’ under supervision of the ‘Animal Welfare Officers’ of Heidelberg University and were approved by the local governing body (Regierungspraesidium Karlsruhe, approval number G16/20). ARRIVE guidelines were followed and sample sizes were calculated as previous experience with G-power analyses.

CHRNA3-EGFP mice (Tg(Chrna3-EGFP)BZ135Gsat/Mmnc) were purchased from the Mutant Mouse Resource & Research Center (MMRRC) and were backcrossed to a C57Bl/6J background. Conditional nociceptor TMEM100 knock-out mice were generated by crossing mice that carry a conditional allele for TMEM100 (B6.Tmem100tm1.1Yjl) ^35^ with SNS-Cre mice C57BL/6-Tg(SCN10A-Cre)1Rkun/Uhg ^36^ (gift from Rohini Kuner). To enable identification of MIAs, these mice were further crossed with CHRNA3-EGFP mice. Furthermore, to identify different nociceptor subclasses for RT-qPCR experiments, Tg(Npy2r-cre)SM19Gsat/Mmucd x B6;129S-Gt(ROSA)26Sortm32(CAG-COP4*H134R/EYFP)Hze/J (Npy2rCre;ChR2-EYFP) mice, in which Aδ-fiber nociceptors express EYFP were used ^6^. Mice were maintained in the Interfaculty Biomedical Facility of Heidelberg University according to institutional guidelines on a 12/12-hour light-dark cycle in an enriched housing environment and had access to food and water ad libitum. For all experiments only adult (age 8-15 weeks) male and female mice were used. Behavioral experiments were conducted at the Interdisciplinary Neurobehavioral Core of Heidelberg University. Prior to the start of experiments animals with the same genetic background and age were randomly assigned to the different experimental groups. To reduce bias investigators were blinded to group identity including treatment (CFA/Saline) and genotype (TMEM100KO/WT).

### HEK293 cell maintenance and transfection

To assess the mechanosensitivity of TMEM100 and of PIEZO2 in the presence and absence of TMEM100, the constructs were (co)-transfected into HEK293 cells using the calcium phosphate method. Cells were grown in DMEM (Thermo Fisher) supplemented with 10% FBS (Thermo Fisher), 2 mM L-glutamine (Thermo Fisher) and penicillin streptomycin (Thermo Fisher, 100 U/mL) at 37ºC and 5% CO2. The day before transfection, cells were seeded on poly-L-lysine treated glass coverslips. For transfection, growth medium was replaced with transfection medium consisting of DMEM, 10% calf serum (Thermo Fisher) and 4 mM L-Glutamine. DNA (0.6 μg/coverslip) was diluted in 100 μl water and after adding CaCl2 2.5M (10 μL per coverslips) the solution was vigorously mixed. Then, 2 x BBS (in mM, 50 HEPES, 280 NaCl, 1.5 Na2HPO4, pH 7.0; 100 μL/coverslip) was added and vortexed. The resulting DNA mix was added to the transfection medium. After 3-4 hours at 37ºC (5% CO2) the transfection medium/DNA mix was replaced with regular HEK293 growth medium. PIEZO2 function was assessed 48 h after transfection.

### Primary DRG cell culture

For primary DRG cultures, mice were sacrificed by placing them in a CO2-filled chamber for 2–4 min followed by cervical dislocation. Lumbar L3 and L4 DRG were collected in Ca2+ and Mg2+-free PBS. DRG were subsequently treated with collagenase IV for 30 minutes (0.5 mg/ml, Sigma-Aldrich, C5138) and with trypsin (0.5 mg/ml, Sigma-Aldrich, T1005) for further 30 minutes, at 37 °C. Digested DRG were washed twice with growth medium [DMEM-F12 (Gibco®, Thermo Fisher Scientific) supplemented with L-glutamine (2 μM, Sigma-Aldrich), glucose (8 mg/ml, Sigma-Aldrich), penicillin (200 U/ml)–streptomycin (200 μg/ml) (Gibco®, Thermo Fisher Scientific), 5 % fetal horse serum (Gibco®, Thermo Fischer Scientific)], triturated using a pipette with filter tips of decreasing diameter (10x up and down with 1000μl filter tip, 10x up and down with 200μl filter tip) and plated in a droplet of growth medium on a glass coverslip precoated with Laminin (GG-12-Laminin coated coverslips, Neuvitro). To allow the dissociated neurons to adhere, coverslips were incubated for 3 hours at 37 °C in a humidified 5 % incubator before being flooded with fresh growth medium. Depending on the experiment neurons were used directly after adding fresh growth medium (see Reverse transcription and quantitative real-time PCR) or after 24h of incubation (see Patch-clamp recordings). For the experiments shown in Fig. 1f – h, primary DRG cultures were transfected using the Amaxa Nucleofector 4D (Lonza) following the manufactures instructions.

### Inflammatory knee pain model

To induce inflammatory knee joint pain the Complete Freund’s Adjuvant (CFA)-induced knee joint monoarthritis model was used as previously described ^53^. In brief, animals were anesthetized in a transparent plexiglass chamber filled with 4% isoflurane in 100% O2 at a flow rate of 1.0 L/min for 3 min. During the procedure anesthesia was maintained using a nosecone delivering a 1.5% Isofluran-O2 mixture while respiratory function was monitored carefully. Adequate anesthesia was confirmed by absence of the pedal reflex (toe pinch). Then, ophthalmic ointment was applied to both eyes to prevent desiccation and the animals were placed in a supine position. Prior to injection of CFA, the left knee was shaved using a commercially available electrical facial hair trimmer, disinfected with povidone-iodine scrub (7.5% solution, Braunol®) and stabilized in a bent position by placing the index finger beneath the knee joint and the thumb above the anterior surface of the ankle joint. The patellar tendon shining through the shaved skin served as visual landmark for the injection. To ensure a precise intraarticular (i.a.) injection the gap inferior to the lower edge of the patella was identified by running a 30 G Insulin syringe horizontally along the knee. To mark the injection level gentle pressure was applied to skin without piercing it leaving behind a horizontal dermal print line. For the injection the needle was lifted vertically at the marked level and inserted at the midline through the patella tendon perpendicular to the tibial axis. Then, the needle was advanced approximately 2-2.5mm without resistance to fully enter the knee joint and 30μl of CFA (1μg/μl) or saline were injected into the joint cavity. After the procedure, the injection site was disinfected and the knee was briefly massaged and mobilized to ensure even distribution of CFA/saline before the animals were returned to their home cages placed on a heating pad for recovery.

### Behavioral testing

All behavioral tests were performed in awake, unrestrained mice by experienced investigators blinded to group identity. Before testing, all animals were habituated to the behavioral test setups at least 3 times over a total of 3 days (1x/d per setup) in the week before starting the behavioral experiment. For the von Frey (vF) and Hargreaves’ test animals were habituated for 1h/setup, and 30 to 60 minutes immediately before each test with the experimenter present in the same room. For the CatWalk XT a habituation session was completed when mice voluntarily crossed the runway 3 times without stopping, turning around, or changing direction (approx. 5 min/animal). On testing days, acclimatization to the CatWalk setup was not necessary. Behavioral assays were always carried out in the same order (CatWalk, vF, Hargreaves) using the same rooms and same test setups at the same time point of the day between 8 a.m - 3 p.m. Prior to knee injection, at least two baseline measurements on two different days were conducted for all behavioral tests. After injections, behavior was evaluated 1, 2, 3, 5 and 7 days post injection (dpi) and then in weekly intervals (14 dpi) for a total of 3 weeks (21dpi).

### Von Frey Test: mechanical sensitivity

Mechanical sensitivity was assessed using the von Frey test as described previously ^54^ 3. Animals were placed in transparent plastic chambers (Modular animal-enclosure; Ugo Basile Srl, Gemonio, Italy) on a 90×38 cm perforated metal shelf (Framed testing surface; Ugo Basile Srl, Gemonio, Italy) that was mounted on a stimulation base. The plantar surface of the animals’ hind paws was perpendicularly stimulated with graded von Frey filaments (Aesthesio® Precision Tactile Sensory Evaluators) of different forces, ranging from 0.07 to 1.4 g without moving the filaments horizontally during application. Each filament was applied 5 times to both the right and left hind paws and the response rate to stimulation in percent (positive response/number of applied stimuli) was used to express mechanical sensitivity. Withdrawal of the stimulated paw was defined as a positive response. Between stimulations of the same hind paw animals had least 1 min break. Prior to knee injection baseline withdrawal frequencies were determined by measuring the withdrawal response rates for all filaments on two different days. After injections, mechanical sensitivity was evaluated 1, 2, 3, 5 and 7 days post injection (dpi) and then in weekly intervals for a total of 3 weeks (21dpi). The 50% withdrawal threshold (WDT) in grams was determined by fitting the response rate vs. von Frey force curves with a Boltzmann sigmoid equation with constant bottom and top constraints equal to 0 and 100, respectively.

### Hargreaves’ test: thermal sensitivity

Thermal sensitivity was assessed according to Hargreaves’ method ^55^ using the Plantar test (Hargreaves Apparatus; Ugo Basile Srl). In brief, withdrawal latency (WDL) in seconds to an infrared (IR) heat beam stimulus applied to the plantar surface of the hind paws was recorded to determine thermal sensitivity. The IR intensity of the radiant heat source was adjusted to obtain baseline WDL between 5 and 7 seconds (IR intensity 50%), with a pre-determined cut-off time of 15 seconds to prevent tissue damage. Each paw was assessed three times and between trials for the same paw, animals had at least an one minute break. To prevent an order effect the paw testing order was chosen randomly. Baseline and post-interventional measurements were conducted at the same time points as described for the vF test.

### CatWalk XT: gait analysis

To quantitatively assess locomotion, the CatWalk XT (version 10.6) gait analysis system (Noldus, Netherlands system) was used. This system consists of an enclosed black corridor (1.3m length) on a glass plate. Inside this glass plate a green LED light is internally reflected. When animals touch the glass plate the light is refracted on the opposite side so that areas of contact become illuminated and detectable. Using the Illuminated Footprints™ technology, videos including the illuminated areas (e.g. paw prints) can be recorded using a high speed color camera (100 frames/s) that is positioned underneath the glass plate. The data is automatically transferred to computer running the CatWalk XT software for further gait analysis. Animals were habituated to the set up as described above (see Behavioral testing). On testing days, animals were placed on one end of the corridor and were allowed to transverse it voluntarily without any external enforcement after setting up the walkway according to the manufacturer’s recommendations. For each mouse on each measurement time point three compliant runs were recorded. A compliant run was defined as a mouse walking across the runway without stopping, turning around, or changing direction and meeting the following pre-determined run criteria: minimum run duration of 0.5s and a maximum run duration of 12s. For all runs the same detection settings were used (camera gain: 16.99, green intensity threshold: 0.10, red ceiling light: 17.7, green walkway light: 16.5). In our analyses we focused on the following gait parameters: stand time in seconds, paw print area in cm^2^, swing speed in cm/s. To better illustrate pain-associated changes in the gait cycle including the stand and swing phase, run data of the left (LH) and right hind paw (RH) are displayed as ratio (LH/RH). For each testing day the ratios (LH/RH) of all three runs per animal were averaged so that the mean of three compliant runs represents the overall result of the animal on that testing day.

### Retrograde labeling

To identify sensory neurons innervating the knee joint retrograde labeling with Fast Blue (FB, #17740-1, Polysciences) was performed. To this end, the same anesthesiological and surgical approach was used as described above (see Inflammatory knee pain model). 2μl of a 4% FB (in saline) solution were injected i.a. in both knee joints using a 10μl Hamilton syringe fitted with 30G needle. After a waiting period of 7d allowing the FB to retrogradely travel to the DRG, animals were further processed depending on the following experiment. To quantify the knee innervating neurons (Fig 3a), animals were sacrificed for microscopy (see Tissue processing and immunochemistry). For electrophysiological and qPCR experiments, the left knee was injected again, but with CFA as described above (see Inflammatory knee pain model) and 3 days thereafter – at the time of maximum pain – the animals were euthanized for primary DRG cultures.

### Patch-clamp recordings

Whole cell patch clamp recordings were made from retrogradely FB-labelled sensory neurons innervating the knee (see Retrograde labeling). To distinguish between MIAs and other peptidergic nociceptors, CHRNA3-EGFP^+^/FB^+^ neurons and small (<30μm) IB4^−^/FB^+^ neurons (see Immunohistochemistry) were recorded, respectively. These two subpopulations account for the majority of nociceptive knee joint afferents (see Fig. 3b). Cells from both WT and TMEM100KO animals after CFA and control treatment were assessed. To this end, 7d after retrograde labeling of the DRG from both knees animals received a second injection into the left knee using CFA or Saline. 3d after this second injection, at the time of maximum CFA-induced pain behavior, the animals were sacrificed. L3 and L4 DRG from the ipsi-(CFA/Saline) and contralateral side were collected separately and cultured for 16-24h (see Primary DRG culture) until used for whole cell patch clamp recordings.

Whole cell patch clamp recordings were made at room temperature (20-24°C) using patch pipettes with a tip resistance of 2-4 MΩ that were pulled (Flaming-Brown puller, Sutter Instruments, Novato, CA, USA) from borosilicate glass capillaries (BF150-86-10, Sutter Instrument). The patch pipettes were filled with a solution consisting of 110 mM KCl, 10 mM NaCl, 1 mM MgCl_2_, 1 mM EGTA, 10 mM HEPES, 2 mM guanosine 5’-triphosphate (GTP) and 2 mM adenosine 5’-triphosphate (ATP) adjusted to pH 7.3 with KOH. The bathing solution contained 140 mM NaCl, 4 mM KCl, 2 mM CaCl2, 1 mM MgCl_2_, 4 mM glucose, 10 mM HEPES and was adjusted to pH 7.4 with NaOH. All recordings were made using an EPC-10 amplifier (HEKA, Lambrecht, Germany) in combination with Patchmaster© and Fitmaster© software (HEKA). Pipette and membrane capacitance were compensated using the auto function of Patchmaster and series resistance was compensated by 70 % to minimize voltage errors. Mechanically activated currents were recorded in the whole-cell patch-clamp configuration. Neurons were clamped to a holding potential of -60 mV and stimulated with a series of mechanical stimuli in 0.8 μm increments with a fire-polished glass pipette (tip diameter 2-3μm) that was positioned at an angle of 45° to the surface of the dish and moved with a velocity of 3 μm/ms by a piezo based micromanipulator called nanomotor© (MM3A, Kleindiek Nanotechnik, Reutlingen, Germany). The evoked whole cell currents were recorded with a sampling frequency of 200 kHz. Mechanotransduction current inactivation was fitted with a single exponential function (C1+C2*exp(–(t–t0)/τ_inact_), where C1 and C2 are constants, t is time and τ_inact_ is the inactivation time constant ^56^.

### Tissue processing and immunochemistry

To quantify retrogradely labelled neurons, DRG were dissected in ice-cooled PBS, fixed with Zamboni’s fixative for 1 h at 4 °C and incubated overnight in 30 % sucrose at 4°C. Then, DRG were embedded in optimum cutting temperature compound (Tissue-Tek™ O.C.T. Compound; Sakura Finetek Germany GmbH, Staufen), cut into 16 μm cryo-sections using a cryostat (Leica CM1950, Leica, Wetzlar, Germany) and mounted onto slides (Microscope Slides SUPERFROST PLUS; Thermo Fisher Scientific, Schwerte, Germany) which were stored at -80°C until used for immunohistochemistry. After drying, sections were treated with 50 mM Glycine in PBS for 20 min, washed twice with 0.2 % Triton X-100 in PBS (0.2% PBST), blocked with 10% normal donkey serum (NDS) and 1% bovine serum albumin (BSA) in 0.2% PBST for 30 min and then incubated with primary antibodies overnight at 4°C. Primary antibodies were diluted in the blocking solution (10% NDS and 1% BSA in 0.2% PBST). Next day, sections were washed 4 × 15 min with 0.2% PBST, subsequently incubated with secondary antibodies for 1 h at room temperature (RT), washed with 0.2% PBST four times (15 min each), dried and coverslipped with FluoroGel (FluoProbes®, Interchim, Montluçon France).

Cultured DRG neurons (see Primary DRG cultures) for electrophysiological and qPCR experiments were counterstained with Alexa Fluor™ -568 conjugated IB4 (2.5μg/ml, Isolectin GS-IB from Griffonia simplicifolia, Alexa Fluor™ 568 Conjugate, Invitrogen™/Thermo Fischer Scientific, I21412) for 10-15 minutes at room temperature to identify different nociceptor subpopulations (CHRNA3-EGFP^+^/FB^+^ and IB4^−^/FB^+^ neurons).

Knees were dissected in cold PBS and fixed with Zambonis fixative overnight. Then, the knee joints were washed in purified water (Milli-Q, Merck KGaA, Darmstadt, Germany) for 3x 30 min before being decalcified by submerging the samples in 10 % EDTA in PBS for 7-10 days (PBS/EDTA was replaced every other day) on a tube roller mixer at 4 °C. After decalcification the samples were washed in PBS for 3 x10 min at RT and cryoprotected in 30% sucrose solution at 4 °C for at least 24h. For the preparation of tissue sections, samples were embedded in optimum cutting temperature compound (Tissue-Tek™ O.C.T. Compound; Sakura Finetek Germany GmbH, Staufen), cut into 25 μm consecutive coronal cryo-sections in arterior-posterior direction and mounted onto microscope slides (SUPERFROST PLUS; Thermo Fisher Scientific, Schwerte, Germany). After drying at RT for 1h, sections were incubated with 50 mM Glycine in PBS for 30 min, washed and blocked 3 × 10 min with 0.5% Tween® 20 in PBS (0.5% PBS-Tw) and then incubated with primary antibodies for 3d at 4°C. Primary antibodies were diluted in a PBS solution containing 1% BSA and 0.3% Triton X-100. Sections were then washed 3 × 10 min with 0.5% PBS-Tw and subsequently incubated for 2 hours at RT with secondary antibodies diluted in PBS with 1% BSA and 0.3% Triton X-100. Finally, the slides were washed with PBS (3 × 10 min), dried and coverslipped using FluoroGel mounting medium with 4,6-diamidino-2-phenylindole (DAPI) counter stain (FluoProbes®, Interchim, Montluçon FRANCE).

Immunostaining images were captured with the Nikon DS-Qi2 camera mounted on a Nikon Ni-E epifluorescence microscope using appropriate filter cubes and identical exposure times for all slides within one experiment.

Silent afferent density (EGFP^+^/CGRP^+^ fibers) was quantified in anatomical regions including FP (Hoffa’s fat pad), LM (lateral meniscus), MM (medial meniscus), LJC (lateral joint capsule), MJC (medial joint capsule) and CL (cruciate ligament) defined according to landmarks as previously described ^57^ using the area fraction tool of NIH ImageJ software (ImageJ 1.53e; Java 1.8.0_172 [64-bit]). In brief, after conversion to 8-bit and background subtraction, image auto local thresholds were set using the Bernsen method. Then, the different immunostaining images (channels) were merged and processed using the image calculator tool to display only double positive (EGFP^+^/CGRP^+^) signals. Finally, the predefined anatomical regions of interest were overlaid and the area fraction determined representing the labelling density of silent afferents per anatomical region in percent. For each animal, at least three photomicrographs per anatomical region were analyzed and averaged. The labelling density per anatomical region for all animals was expressed as mean ± SEM. For illustration purposes (Fig. 2a) representative images of coronal 100 μm knee sections (Cryostat) were acquired and stitched using a Leica SP8 Confocal microscopy platform equipped with a Lasx 3.5. Laser, detector powers were optimized for the combination of antibodies.

### Antibodies

The following primary antibodies were used: rat anti-GFP (Nacalai tesque, #04404-84, 1:3000), rabbit anti-CGRP (ImmunoStar,#24112, 1:200), Isolectin GS-IB from Griffonia simplicifolia Alexa Fluor™ 568 Conjugate (2.5μg/ml, Invitrogen™/Thermo Fisher Scientific, #I21412), Isolectin GS-IB from Griffonia simplicifolia Alexa Fluor™ 647 Conjugate (2.5μg/ml, Invitrogen™/Thermo Fisher Scientific, #I32450) and rabbit anti-dsRed (1:1000; Takara). The following corresponding Alexa Fluor™ conjugated secondary antibodies (1:750; Thermo Fisher Scientific) were used: Alexa Fluor 488 conjugated donkey anti-Rat IgG (Thermo Fisher Scientific,#A48269), Alexa 594 conjugated donkey anti-Rabbit IgG (Thermo Fisher Scientific, #A32754).

### Reverse transcription and quantitative real-time PCR

To compare NGF-induced changes in mRNA expression levels of TMEM100 in different nociceptor subclasses (Fig. 1e), primary L3-4 DRG neurons from WT mice were cultured in the absence and presence of NGF (50 ng/ml) for 24h before cell collection. To compare mRNA expression levels of TMEM100 in treated (CFA) and control (saline) knee innervating nociceptors at the time of maximum pain (3 dpi), CHRNA3-EGFP^+^/FB^+^ and IB4^−^/FB^+^ neurons from the ipsi-(CFA) and contralateral (saline) side were collected from acute primary L3-4 DRG cultures of WT mice immediately after adding fresh growth medium without further incubation. In both approaches cultures were counterstained with Alexa Fluor™ -568 conjugated IB4 (2.5μg/ml, Isolectin GS-IB from Griffonia simplicifolia, Alexa Fluor™ 568 Conjugate, Invitrogen, I21412) for 10-15 minutes at room temperature to enable the identification of different nociceptor subpopulations.

Samples (20 cells per subpopulation and condition) were manually collected using a fire polished pipette with a tip diameter of ∼25 μm pulled (Flaming-Brown puller, Sutter Instruments, Novato, CA, USA) from borosilicate glass capillaries (BF150-86-10, Sutter Instrument) that were filled with 2μl of picking buffer [1μL RNAse inhibitor (Takara #2313A) in 49 μL PBS]. After aspirating 20 cells per sample [NGF ±: CHRNA3^+^, IB4^−^, IB4^+^, Aδ-nociceptors (see Fig. 1e); CFA/Saline: CHRNA3-EGFP^+^/FB^+^, IB4^−^/FB^+^ (see Fig. 3c)] the pipette was immediately shock frozen in liquid Nitrogen and the cells were expelled into an RNAse free tube filled with 8 μL of picking buffer. Directly thereafter, the tubes were stored at -80°C until further processing. For each gene 4 to 9 samples (1 sample per subpopulation and condition per mouse) were collected. Cell populations were identified and picked using a 20x magnification and appropriate filter cubes in the Zeiss Axio Observer A1 microscope (Carl Zeiss). Cell lysis and reverse transcription with cDNA synthesis was carried out directly on the sample using the Power SYBR® Green Cells-to-CT™ Kit (Thermo Fischer Scientific, #4402953) following the manufacturer’s instructions. qPCR reactions were set up using FastStart Essential DNA Green Master (Roche, #06402712001) according to the manufacturer’s guidelines. Per reaction (20 μl reaction volume) 4 μl of the obtained cDNA as template was added to 10 μl SYBR Green PCR Master Mix, 4 μl nuclease-free H2O and the following forward (FW) and reverse (RV) primer pairs (1μl each of a 5μM dilution, final concentration: 250nM): GAPDH-FWD 5’-GCATGGCCTTCCGTGTTC-3’; GAPDH-REV 5’-GTAGCCCAAGATGCCCTTCA-3’; TMEM100-FWD 5’-GAAAAACCCCAAGAGGGAAG-3’; TMEM100-REV 5’-ATGGAACCATGGGAATTGAA-3’. qPCR reactions were performed in a LightCycler 96 (Roche) with a thermal cycler profile as follows: 10 min preincubation step at 95°C followed by 40 cycles of PCR with a 10 second denaturing cycle at 95°C, followed by 10 seconds of annealing at 60ºC and 10 seconds extension at 72°C. Mean ± SEM expression levels of TMEM100 normalized to the expression levels of the housekeeping gene GAPDH were compared in the different nociceptor subclasses cultured in absence and presence of NGF (Fig. 1e). CFA-induced changes in mRNA expression levels of TMEM100 in CHRNA3-EGFP^+^/FB^+^ and IB4^−^/FB^+^ neurons compared to contralateral control neurons were analyzed using the ΔΔCt method (Fig. 3c).

### RNA Sequencing

For RNAseq, CHRNA3-EGFP^+^ samples (20 cells per sample and condition) of 3 WT mice were processed, cultured (±NGF for 24h) and collected as described above (see Reverse transcription and quantitative real-time PCR). The SmartSeq2 protocol published by Picelli et al. (2014) was used to process cell lysates to reverse transcription and library preparation was performed using the Nextera DNA Sample Preparation kit (Illumina) following the manufacturer’s instructions. Libraries were sequenced with Illumina HiSeq 2000. Sequencing reads were mapped to GRCm38 mouse reference genome and differential gene expression analysis was performed using the BioJupies platform ^59^ with default parameters. Next generation RNA-sequencing raw data (FASTQ files) have been deposited in the Gene Expression Omnibus (GEO) under accession number GSE199580 and are publicly available as of the date of publication.

### Calcium Imaging

To examine the responsiveness of FB-labelled neurons to the TRPA1 agonist allylisothiocyanate (AITC, Sigma-Aldrich) Calbryte-590 (Calbryte™ 590 AM, AAT Bioquest) Ca^2+^-imaging was performed. CHRNA3-EGFP^+^/FB^+^ neurons and small (<30μm) IB4^−^/FB^+^ neurons from both WT and TMEM100KO animals after CFA and control treatment were obtained as described above (see Patch clamp recordings), counterstainded with Alexa Fluor™ -647 conjugated IB4 (2.5μg/ml, Isolectin GS-IB from Griffonia simplicifolia, Alexa Fluor™ 647 Conjugate, Invitrogen™/Thermo Fischer Scientific, I32450) for 10-15 min at room temperature, washed with extracellular buffer (140 mM NaCl, 4 mM KCl, 2 mM CaCl_2_, 1 mM MgCl_2_, 4 mM glucose, 10 mM HEPES and was adjusted to pH 7.4 with NaOH) and then incubated with the Ca^2+^ indicator Calbryte-590 (5μM diluted in ECB from a 5 mM stock solution in DMSO) for 30 min at 37 °C. Coverslips with loaded cells were then washed with ECB, mounted onto a perfusion chamber and superfused with ECB using a constant laminar flow provided through an 8-channel valve controlled gravity-driven perfusion system (VC3-8xG, ALA Scientific Instruments) and a peristaltic pump. A manifold system with 8 inlet ports fitted to a silicon tube bath inlet whose end was positioned at the outer edge of the coverslip without interfering the visual field was used to provide immediate release of ECB and chemical agents into the superfusion chamber. This system enabled minimal dead volume and air bubbles in the lines. Tubes were identical for each input line. All experiments were conducted at room temperature (23 ± 1 °C). Fluorescence images were captured with a Hamamatsu ORCO-Flash4.0 camera at 2 Hz under an inverted Zeiss Axio Observer A1 microscope equipped with a LED light source (CoolLED pE-340fura). ZEN 2 pro software (Carl Zeiss Microscopy GmbH) was employed to detect and analyze intracellular calcium changes throughout the experiment. During imaging the following protocol was applied. After establishing a 30 s baseline with ECB (0-30 s), neurons were challenged with AITC (10 μM) for 60 s (31-90 s) followed by a wash out period of 270 s (91-360 s). At the end of the protocol, 100 mM KCl was applied for 30 s (361-390) to depolarize neurons in order to identify viable neurons in contrast to non-neuronal cells or non-functioning neurons. KCl application was followed by a last wash out period with ECB for 30 s (391-420 s). Neuronal viability was defined as a >20% increase of fluorescence intensity from the mean intensity 20 s pre-KCl application (330 -350s).

Analysis was conducted by extracting mean intensity values of neurons (CHRNA3-EGFP^+^/FB^+^ neurons and IB4^−^/FB^+^) after background subtraction from manually drawn regions of interests (ROIs including background ROI) in the ZEN 2 pro software (Carl Zeiss Microscopy GmbH). These values were then transferred into a custom-made Microsoft Excel® template to compute the proportion of neurons responding to AITC under different conditions (CFA/saline; WT/TMEM100KO). In brief, fluorescence is shown as ΔF/F with ΔF = F1 – F (F1 =mean intensity of image, F = mean intensity of baseline fluorescence from 0-20 s). To express percent changes in fluorescence intensity, ΔF was normalized to F and multiplied by 100 (%). Cells responding with an increase > 5 % of fluorescence intensity from baseline to AITC application were counted as AITC responders. Cells not crossing the KCl threshold were excluded from the analysis.

### Ex-vivo skin-nerve preparation

To examine peripheral sensitization, we directly measured the mechanosensitivity of C-fiber and Aδ-fiber nociceptors in the tibial nerve by recording mechanically evoked action potentials from single nerve fibers in an ex-vivo skin-nerve preparation. To this end, WT and TMEM100KO mice were sacrificed 3d after CFA/saline injection by placing them in a CO2-filled chamber for 2–4 min followed by cervical dislocation. After dissection, the glabrous skin of the hind limb was placed with the corium side up in a heated (32°C) organ bath chamber that was perfused with synthetic interstitial fluid (SIF buffer) consisting of 108 mM NaCl, 3.5 mM KCl, 0.7 mM MgSO_4_, 26 mM NaHCO_3_, 1.7 mM Na H_2_PO_4_, 1.5 mM CaCl_2_, 9.5 mM sodium gluconate, 5.5 mM glucose and 7.5 mM sucrose at a pH of 7.4. The tibial nerve was attached in an adjacent chamber for fiber teasing and single-unit recording. As previously described ^6^, single units were isolated using a mechanical search stimulus applied with a glass rod and classified by conduction velocity, von Frey hair thresholds and adaptation properties to suprathreshold stimuli. A cylindrical metal rod (diameter 1 mm) that was driven by a nanomotor® (MM2A-LS, 914 Kleindiek Nanotechnik GmbH, Germany) coupled to a force measurement system (FMS-LS, Kleindiek Nanotechnik GmbH, Germany) was used to apply mechanical ramp-and-hold stimuli. The mechanical thresholds of single units were determined by mechanically stimulating the most sensitive spot of the receptive fields using von Frey filaments (Aesthesio® Precision Tactile Sensory Evaluators). The force exerted by the weakest von Frey filament that was sufficient to evoke an action potential was considered as the mechanical threshold. The raw electrophysiological data was amplified with an AC coupled differential amplifier (Neurolog NL104 AC), filtered with a notch filter (Neurolog NL125-6), converted into a digital signal with a PowerLab SP4 (ADInstruments) and recorded at a sampling frequency of 20 kHz using LabChart 7.1 (ADInstruments).

### AAV-PHP.S production

AAV-PHP.S viral particles were produced using a modified protocol based on established procedures by Gradinaru and colleagues ^60^. Briefly, AAV-293 cells (Agilent, 240073) were seeded on 150mm dishes and transfected using polyethylenimine (Polysciences, 23966) with four plasmids: a pAAV of interest (AAV-CAG-dsRedExpress2 or AAV-CAG-TMEM100-IRES-dsRedExpress2, both with AAV2 ITRs), pAdDeltaF6 (Helper, Addgene #112867), pUCmini-iCAP-PHP.S (Addgene #103006) and a mutated pUCmini-iCAP-PHP.S having a 6xHis tag on the VP3 capsid protein ^61^, with 1:2:2:2 ratio respectively. Cell culture medium was changed at 48 and 120 h post-transfection and supernatant containing viral particles was centrifuged at 1690 g for 10 minutes. Supernatant medium was filtered (0.2μm) and diluted in PBS. Cell pellet and filtered medium were stored at 4°C. At 120h post transfection, pelleted cells and the ones in the dishes were lysed and incubated at 37°C for 1 hour with the specific PBS buffer containing: MgCl_2_ 6mM, Triton X-100 0.4%, RNAse A 6μg/ml (Roche, #10109169001), DENARASE 250U/μl (c-LEcta, #20804). Lysed cells were collected, diluted in PBS and centrifuged at 2300 g for 10 minutes. Supernatant from the cell lysate and the filtered medium were incubated separately with equilibrated Ni-sepharose excel histidine-tagged protein purification resin (Cytiva, #17371202) for at least 2 hours at room temperature with gentle mixing. Filtered medium and cell lysate were carefully loaded through a gravity flow chromatography column with a 30μm filter (Econo-pac, Bio Rad, #7321010). Beads were washed with 80ml of washing buffer (20 mM imidazole in PBS, pH 7.4) and viral particles were then eluted in 50ml of elution buffer (500 mM imidazole in PBS, pH 7.4). Buffer exchange and concentration was done with Vivaspin 20 ultrafiltration unit having a 1.000.000 molecular weight cut-off (Sartorius). Viral particles were washed and resuspended in PBS and titered using quantitative PCR with primers targeting WPRE element.

### Statistics

Unless otherwise stated, all data are expressed as means ± s.e.m. All statistical analyses were performed with Microsoft Excel and Prism 9.0 (Graphpad). Data distribution was systematically evaluated using D’Agostino-Pearson test and parametric or non-parametric tests were chosen accordingly. The statistical tests that were used, the exact P-values and information about the number of independent biological replicates are provided in the display items or the corresponding figure legends. Symbols on graphs (* or #) indicate standard P-value range: *, P < 0.05; **, P < 0.01; ***, P < 0.001 and ns (not significant) P > 0.05. Additional information about the statistical tests is provided in Data S1 in the Supplementary Information.

## Supporting information

Source data and statistical information

## Data and code availability

All data supporting the findings of this study are available within the article and its supplementary information files. All RNA sequencing dataset generated in this study are deposited in the Gene Expression Omnibus under accession number GSE199580. Plasmids generated in this study are available from the corresponding author upon request. Statistical source data are provided with this paper. A reporting summary for this article is available as a Supplementary Information file.

## Acknowledgements

We thank Ms. Anke Niemann and Ms. Claudia Lüchau for technical support. We also acknowledge support from Dr. Claudia Pitzer from the Interdisciplinary Behavioural Core (INBC) at Heidelberg University. This work was funded by the Deutsche Forschungsgemeinschaft grant SFB1158/A01 to S.G.L. and a fellowship from the Physician Scientist Program of the Medical Faculty of Heidelberg University awarded to T.A.N.

## Author Contributions

T.A.N., N.W., C.V., P.A., I.S., S.B., J.V., V.P., C.M., N.Z., F.J.T. and S.G.L. performed the experiments and analyzed data. G.R.L., Y.J.L. and P.A.H. provided material. T.A.N. and S.G.L. designed and supervised the experiments and wrote the manuscript.

