## Supplementary information for "The molecular mechanism and physiological role of silent nociceptor activation"

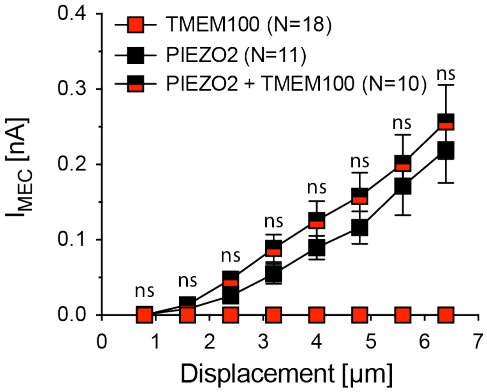


**Supplementary Figure 1, TMEM100 does not produce mechanotransduction currents or modulates PIEZO2-mediated currents, related to Figure 1**
Comparison of the mean ± SEM peak current amplitudes of mechanically-evoked currents in HEK293 cells expressing PIEZO2 (black squares), PIEZO2+TMEM100 (half-filled square) or TMEM100 (red squares) alone. N-numbers are indicated in the graph legend. PIEZO2 and PIEZO2+TMEM100 currents were compared using multiple Mann-Whitney tests (ns, not significant).
Source data and statistical information are provided as a Source Data file.

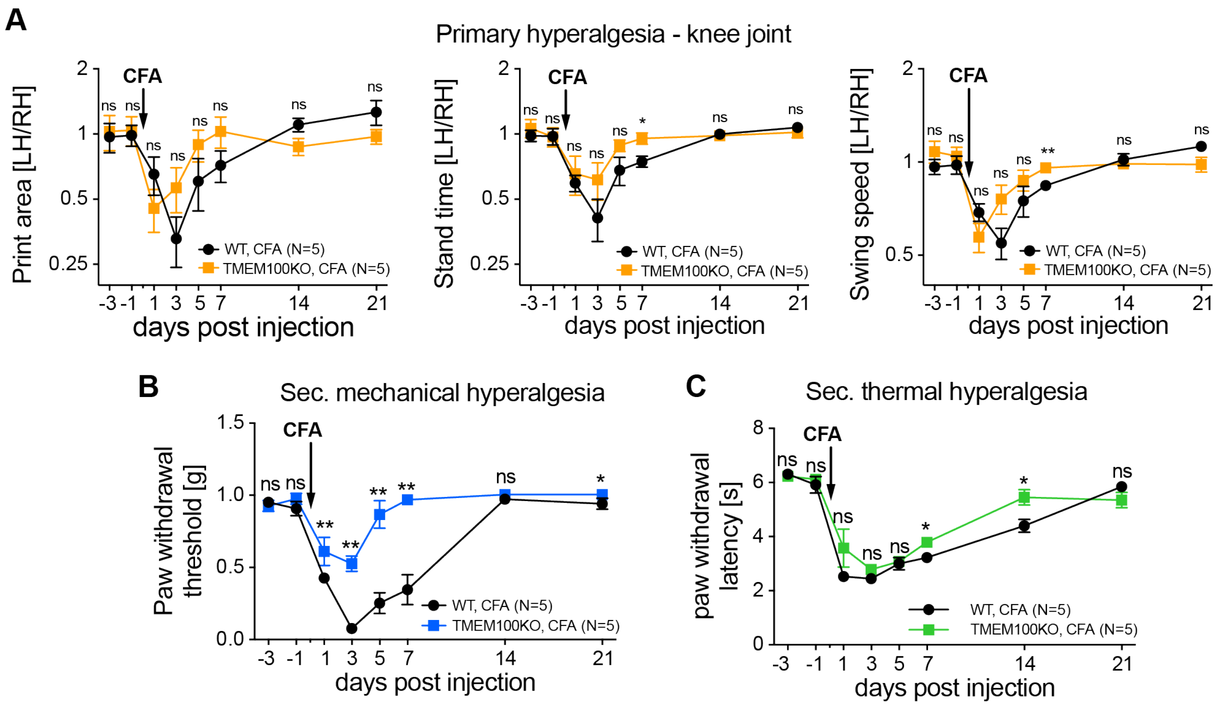


**Supplementary Figure 2, Female TMEM100KO mice exhibit the same pain phenotype as male mice, related to Figure 4**

**(A)**, Comparison of the time courses of changes in the ratios (left/right hind paw) of print area (left), stand time (middle) and leg swing speed (right) of female CFA injected WT mice (black circles), female CFA injected TMEM100KO mice (orange squares). Symbols represent means ± SEM, N-numbers are provided in the graph legends and ratios at different time points were compared using multiple Mann-Whitney tests (ns, not significant; *, $P<0.05$; **, $P<0.01$).

**(B)**, Comparison of the time courses of changes in the mechanical paw withdrawal thresholds of female CFA injected WT mice (black circles), female CFA injected TMEM100KO mice (blue squares). Symbols represent means ± SEM, N-numbers are provided in the graph legends and ratios at different time points were compared using multiple Mann-Whitney tests (ns, not significant; *, $P<0.05$; **, $P<0.01$).

**(C)**, Comparison of the time courses of changes in the thermal paw withdrawal latencies determined with the Hargreaves test of female CFA injected WT mice (black circles), female CFA injected TMEM100KO mice (blue squares). Symbols represent means ± SEM, N-numbers are provided in the graph legends and ratios at different time points were compared using multiple Mann-Whitney tests (ns, not significant; *, $P<0.05$; **, $P<0.01$).

Source data and statistical information are provided as a Source Data file.

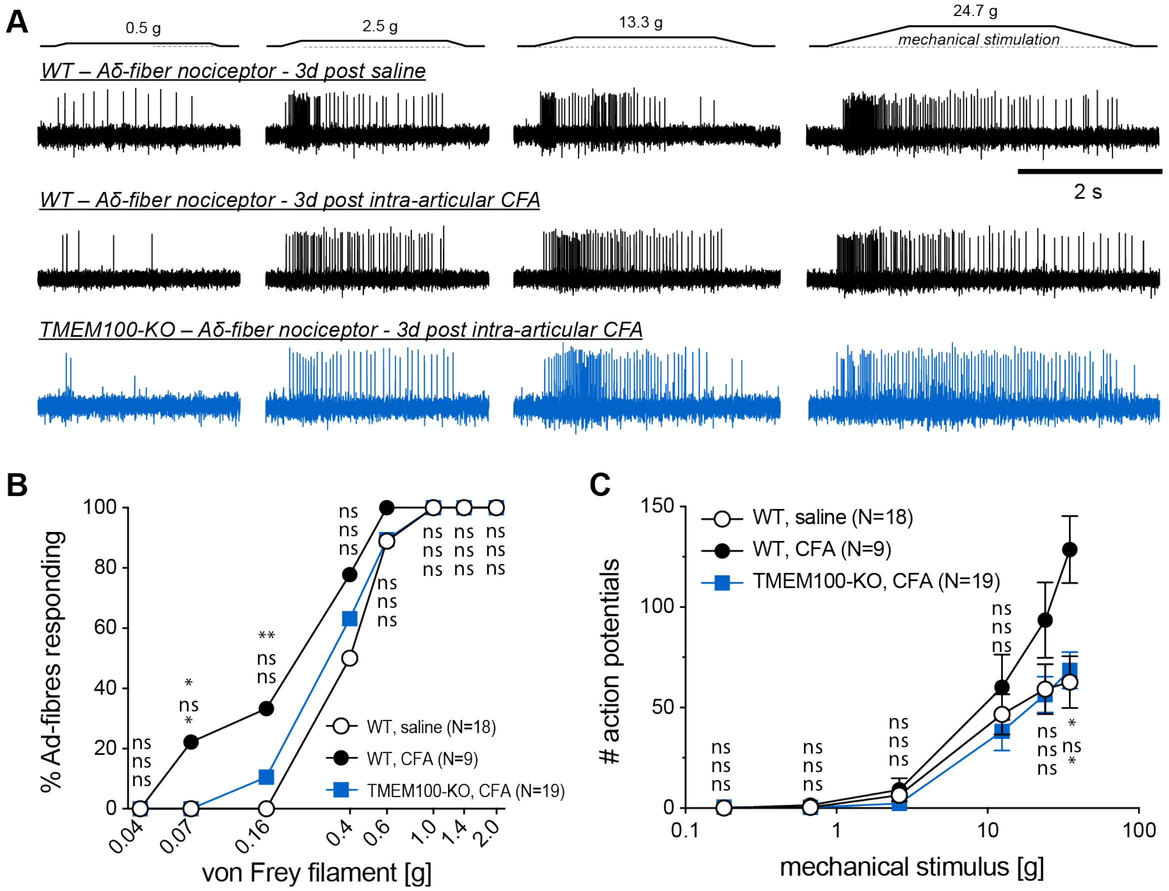


**Supplementary Figure 3, Sensitization of cutaneous Aδ-fiber nociceptors by intraarticular CFA, related to Figure 6**

**(A)**, Example traces of mechanically-evoked action potentials recorded from single nerve Aδ-fibers from the tibial nerve of WT mice 3 days post saline (top), WT mice 3 dpi CFA (middle) and TMEM100KO mice 3 dpi CFA (bottom).

**(B)**, Comparison of the proportions of Aδ-fiber nociceptors that respond to mechanical stimulation with the indicated von Frey filaments. The proportions were compared pairwise using the Chi-square test. P-values are provided next to the symbols in the graph and refer to WT-saline vs. WT-CFA (top), WT-saline vs. TMEM100KO-CFA (middle) and WT-CFA vs. TMEM100KO (bottom). N-numbers are indicated in the graph legend.

**(C)**, Comparison of the firing rates of Aδ-fibers evoked by a series of ramp-and-hold stimuli with increasing amplitudes that exerted the indicated force to the receptive fields. Symbols represent the mean ± SEM numbers of action potentials, which were compared using multiple Mann-Whitney tests. P-values (ns, not significant; *, P<0.05) are provided next to the symbols in the same order as in (**B**).

Source data and statistical information are provided as a Source Data file.

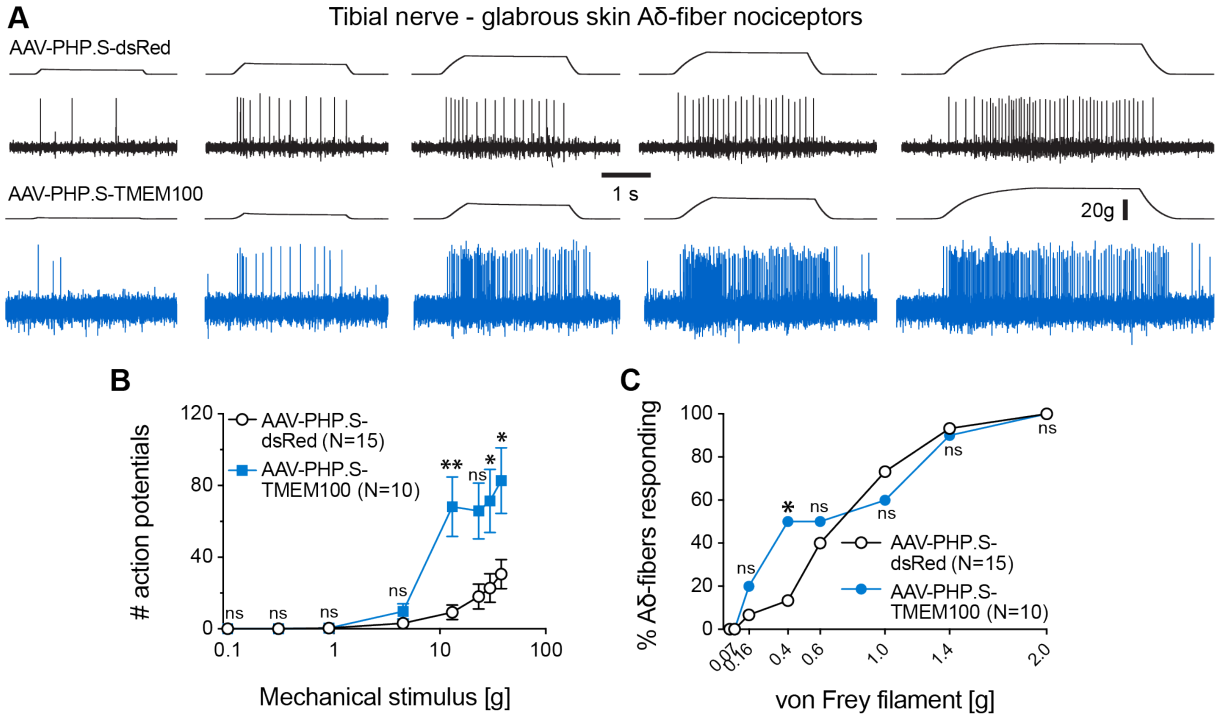


**Supplementary Figure 4, Sensitization of cutaneous Aδ-fiber nociceptors by TMEM100 overexpression in articular afferents, related to Figure 7**

**(A)**, Example traces of mechanically-evoked action potentials recorded from cutaneous Aδ-fiber nociceptors in the tibial nerve from control mice (top, AAV-PHP.S-dsRed) and from mice that overexpress TMEM100 in articular afferents (bottom, AAV-PHP.S-TMEM100-Ires-dsRed).

**(B)**, Comparison of the firing rates evoked by ramp-and-hold stimuli that exerted the indicated force to the receptive fields. Symbols represent means ± SEM numbers of action potentials, which were compared using multiple Mann-Whitney tests (ns, not significant; *, P<0.05; **, P<0.01).

**(C)**, Comparison of the proportions of C-fiber nociceptors that respond to mechanical stimulation with the indicated von Frey filaments. The proportions were compared pairwise using the Chi-square test (ns, not significant; *, P<0.05).

Source data and statistical information are provided as a Source Data file.
